# Inferring neural signalling directionality from undirected structural connectomes

**DOI:** 10.1101/573071

**Authors:** Caio Seguin, Adeel Razi, Andrew Zalesky

## Abstract

Neural information flow is inherently directional. To date, investigation of directional communication in the human structural connectome has been precluded by the inability of non-invasive neuroimaging methods to resolve axonal directionality. Here, we demonstrate that decentralized measures of network communication, applied to the undirected topology and geometry of brain networks, can predict putative directions of large-scale neural signalling. We propose the concept of send-receive communication asymmetry to characterize cortical regions as senders, receivers or neutral, based on differences between their incoming and outgoing communication efficiencies. Our results reveal a send-receive cortical hierarchy that recapitulates established organizational gradients differentiating sensory-motor and multimodal areas. We find that send-receive asymmetries are significantly associated with the directionality of effective connectivity derived from spectral dynamic causal modeling. Finally, using fruit fly, mouse and macaque connectomes, we provide further evidence suggesting that directionality of neural signalling is significantly encoded in the undirected architecture of nervous systems.

## INTRODUCTION

Understanding how the structural substrate of connectomes [1, 2] gives rise to the rich functional dynamics observed in nervous systems is a major goal in neuroscience [3–6]. Anatomical connectivity constrains and facilitates neural information transfer, which in turn gives rise to synchronization (i.e., functional connectivity) between neural elements. Therefore, knowledge of how neural signals are communicated in nervous systems can establish a bridge between structural and functional descriptions of brain networks [7–10].

While information can be directly communicated between anatomically connected neural elements, polysynaptic communication is needed for structurally unconnected elements. Network communication models describe a propagation strategy that delineates the signalling pathways utilized to transfer information between network nodes. In turn, a network communication measure quantifies the *communication efficiency* along the identified pathways from a graph-theoretic standpoint. Efficient communication pathways are generally short, traverse few synapses and comprise strong and reliable connections [11].

Several network models of polysynaptic communication have been proposed [9]. Shortest paths routing is the most ubiquitous model [12–14], which proposes that communication occurs via optimally efficient routes. However, the identification of shortest paths mandates global knowledge of network topology [8, 9, 15, 16]. This requirement is unlikely to be met in biological systems, in which individual elements (e.g., neurons or brain regions) do not possess information on all connections comprising the network. This has motivated research on decentralized models that capitalize on local knowledge of network properties to facilitate signalling. Examples include navigation [15, 17, 18], spreading dynamics [19, 20], and diffusion processes [21–24].

Many decentralized communication models are asymmetric [8, 15, 16, 19], meaning that sending information from region *i* to region *j* can be performed more efficiently than sending information from region *j* to region *i*. We coin the term *send-receive communication asymmetry*, or simply *send-receive asymmetry* to describe this property. Importantly, the interplay between decentralized communication, network topology and possibly geometry can result in communication asymmetry in undirected networks (Fig. 1). This provides an opportunity to infer putative directions of information flow from current descriptions of the human structural connectome, for which knowledge about the directionality of individual connections is unknown due to inherent limitations of *in vivo* diffusion imaging. Therefore, decentralized network communication measures may help bridge the gap between our symmetric understanding of human connectome structure and the ample evidence for its asymmetric functional dynamics [25–27].

**FIG. 1.**
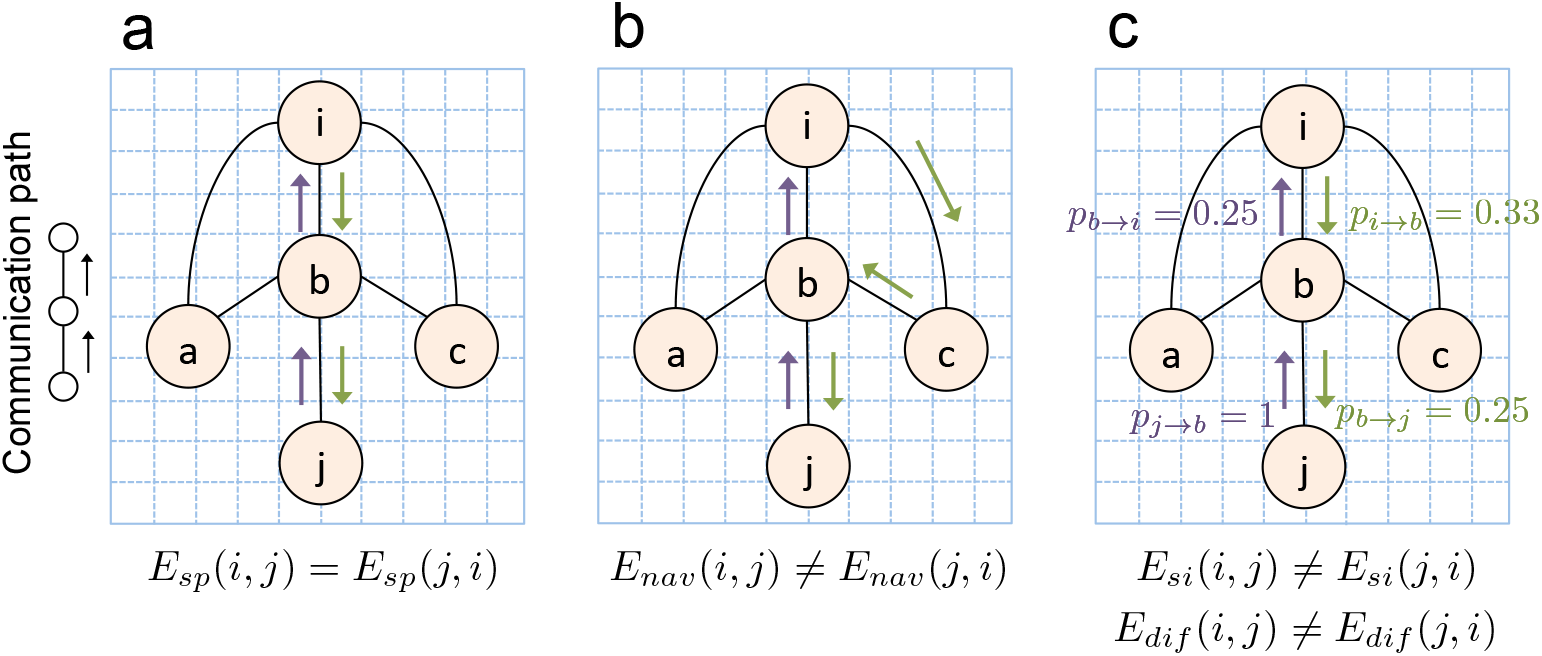
Illustrative examples of send-receive communication asymmetry. The toy network is spatially-embedded, unweighted and undirected. Communication efficiency from node *i* to *j* under measure *x* ∈ {*sp,nav,si,dif*} is denoted *E_x_*(*i, j*). Shortest path and navigation efficiencies are computed as the inverse of the number of connections comprising shortest and navigation paths, respectively. Diffusion efficiency relates to how quickly, on average, a random walker can travel between two nodes, while search information relates to the probability that a random walker will travel between two nodes via the shortest path linking them. The path identified under each communication model is designated with green (*i* → *j*) and mauve (*j* → *i*) arrows. Send-receive communication asymmetry refers to *E_x_*(*i, j*) ≠ *E_x_*(*j, i*). **(a)** Shortest path efficiency is always symmetric in undirected networks, and thus *E_sp_*(*i, j*)=*E_sp_*(*j, i*). **(b)** Navigation routes information by progressing to the next directly connected node that is closest in distance to the target node. This results in the *i-c-b-j* and *j-b-i* navigation paths, with respective efficiency *E_nav_*(*i,j*) = 0.33 and *E_nav_*(*j, i*) = 0.5. Hence, navigation is more efficient from node *j* to node *i*. **(c)** Arrows denote the symmetric shortest paths between *i* and *j*. Arrows are annotated with the probabilities that a random walker will traverse their respective connections based on node degree (e.g., each of the 3 connections of node *i* has approximately 0.33 probability to be traversed by a random walker leaving *i*). We have *E_si_*(*i,j*) ∝ 0.33 × 0.25 = 0.0825 and *E_si_*(*j,i*) ∝ 1 × 0.25 = 0.25. Hence, a random walker has higher probability of travelling via the shortest path in the *j* → *i* direction, characterizing search information asymmetry between *i* and *j*. Similarly, on average, a random walker is expected to visit fewer nodes travelling from *i* to *j* than from *j* to *i*. Hence, *E_dif_*(*j, i*) > *E_dif_*(*i, j*), characterizing diffusion efficiency asymmetry.

We provide multiple lines of evidence suggesting that decentralized communication measures, applied to undirected brain networks, can provide new insight into the directionality of neural information flow. We begin by classifying individual cortical regions and subsystems as senders (biased towards the efficiency of outgoing paths), neutral (symmetric communication efficiency) and receivers (biased towards the efficiency of incoming paths). We demonstrate that regional variation in send-receive asymmetry recapitulates established hierarchies of cortical organization. Next, we analyze pairwise send-receive asymmetries between cortical subsystems, providing multi-scale maps of communication directionality in the human cortex. Crucially, we validate these maps by showing a significant association between send-receive asymmetry and directionality of effective connectivity derived from dynamic causal modeling (DCM) applied to resting-state functional magnetic resonance imaging (fMRI). These results suggest that the undirected topology of the human connectome imposes constraints on neural signalling directionality. We further investigate this notion by examining fly, mouse and macaque connectomes. We leverage the presence of directed connections in these brain networks to provide additional evidence that neural signalling directionality is not exclusively contingent on directed axonal projections, being partially shaped by the undirected organization of nervous systems.

## RESULTS

### Measures of send-receive communication asymmetry

We investigated three asymmetric network communication measures: i) navigation efficiency, ii) diffusion efficiency and iii) search information (*Materials and Methods, Network communication measures*). Briefly, navigation efficiency [15] relates to the length of paths identified by navigation or greedy routing [17, 28], with higher values of efficiency indicating faster and more reliable communication between nodes. Diffusion efficiency [23] quantifies how many intermediate regions (synapses), on average, a naive random walker needs to traverse to reach a desired destination region. Finally, search information is related to the probability that a random walker will travel from one region to another via the shortest path between them [8, 29], quantifying the extent to which efficient routes are hidden in the network topology. Together, these measures are representative of different conceptualizations of decentralized network communication [9, 16], from single-path routing via geometric navigation to diffusive signalling unfolding along multiple network fronts.

Communication asymmetry is introduced by the decentralized character of certain network communication models (Fig. 1). Consider the flow of information from one region, termed the *source node*, to another region, termed the *target node*. If this source-target pair is not directly connected, information must flow via a polysynaptic path that traverses one or more intermediate nodes. Decisions on how signals are propagated through the connectome depend on the local topology around each node. Since source and target nodes occupy potentially different vicinities, communication may happen through distinct paths, and thus with different efficiency, depending on the direction of information flow. In contrast, centralized communication models such as shortest path routing always yield symmetric paths in undirected networks.

We use *C* ∈ ℝ^*N* × *N* × *K*^ to denote a set of communication matrices for *K* individuals, where *C*(*i, j, k*) denotes the communication efficiency from node *i* to node *j* for individual *k*, under an arbitrary communication measure (Fig. 2a). The difference in communication efficiency for opposing directions of information flow between *i* and *j* is given by Δ(*i, j, k*)=*C*(*i, j, k*) − *C*(*j, i, k*). We perform a one-sample t-test to determine whether the mean of the distribution Δ(*i, j, k* = 1…*K*) is significantly different to 0 (Fig. 2c). This yields a t-statistic, termed *A*(*i, j*), which quantifies the extent of communication asymmetry between *i* and *j*. In particular, if *A*(*i, j*) is significant and greater than zero, we conclude that communication can occur more efficiently from node *i* to node *j*, rather than from node *j* to node *i*. Note that *A*(*i, j*)=−*A*(*j, i*), and thus if *A*(*j, i*) is significantly less than zero, we reach the same conclusion. Repeating this test independently for all pairs of nodes yields the communication asymmetry matrix *A*, for which values are symmetric about the main diagonal, but with opposite signs.

**FIG. 2.**
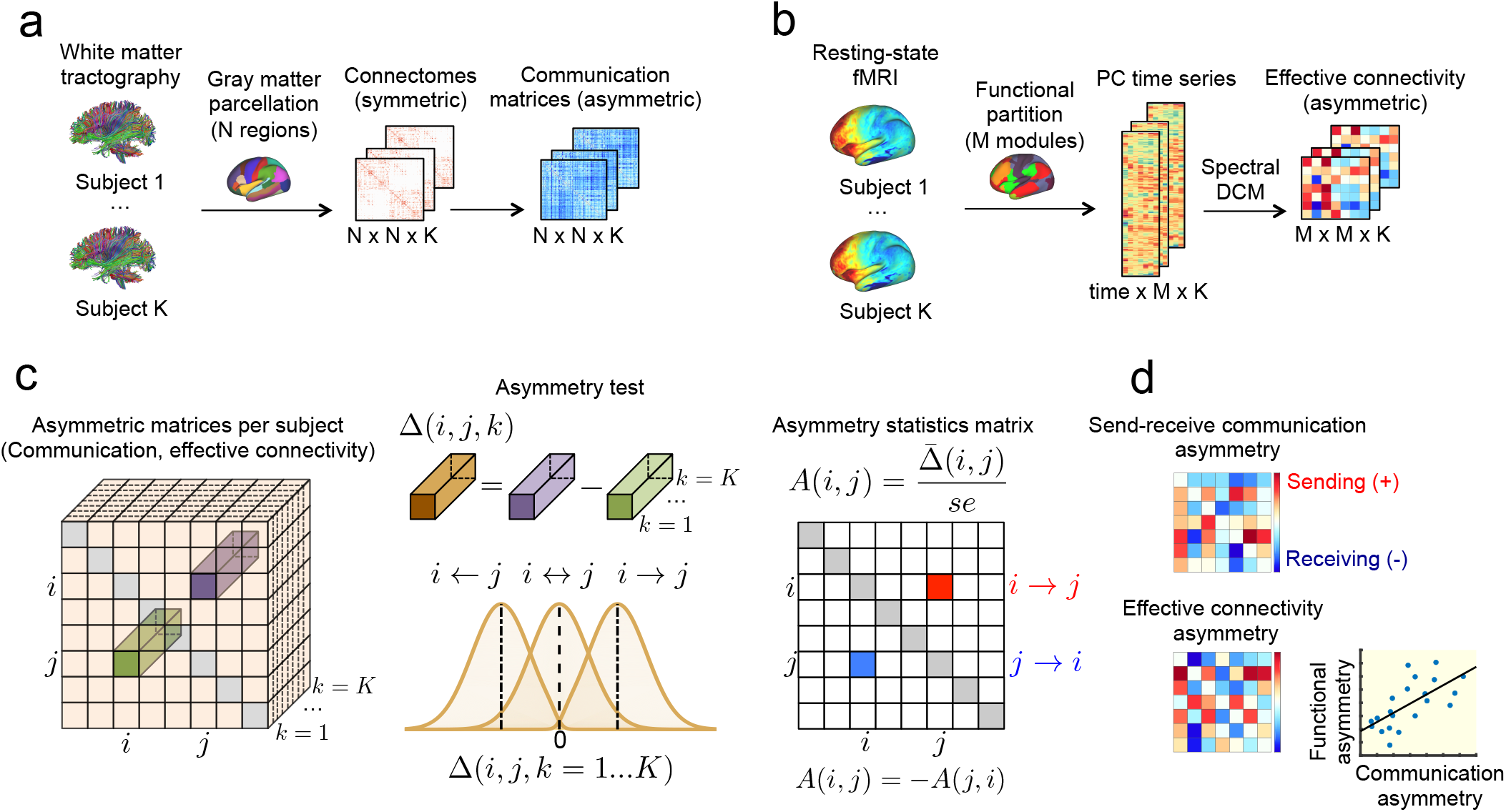
Methodology overview. **(a)** White matter tractography applied to diffusion MRI data for *K* = 200 adults participating in the HCP was used to map undirected (i.e., symmetric) weighted adjacency matrices representing the structural connectivity between *N* =256, 350, 512 cortical regions. Navigation efficiency, diffusion efficiency and search information were computed between every pair of regions to generate asymmetric communication matrices. **(b)** Resting-state fMRI data for the same HCP participants was used to compute principal component (PC) time series summarizing the functional activity of *M* =7, 17, 22 cortical subsystems. For each individual, effective connectivity between cortical subsystems was computed using spectral DCM. **(c)** Schematic of the communication asymmetry test. First, for a pair of nodes *i* and *j*, the difference in communication efficiency between the *i* → *j* and *j* ← *i* directions was computed. Performing this for *K* individuals yielded the distribution Δ(*i, j, k* = 1…*K*). Communication asymmetry was assessed by performing a one-sample t-test to determine whether the mean of this distribution is significantly different to 0, with *A*(*i, j*) defined as the resulting matrix of t-statistics. **(d)** The asymmetry test was applied to compute *M* × *M* matrices of communication and effective connectivity send-receive asymmetries between modules. We sought to test for correlations across the corresponding elements of these two matrices.

The above measure is specific to pairs of nodes. We use a variation of this test to compute regional send-receive communication asymmetry by taking into account all outgoing and incoming communication paths of a given node (*Materials and Methods, Communication asymmetry test* and Supplementary Fig. 1). Regions that show a significantly higher efficiency of outgoing (incoming) communication are classified as putative senders (receivers), while nodes that do not favour a direction of information flow are classified as neutral.

### Senders and receivers of the human connectome

Whole-brain white matter tractography was applied to high-resolution diffusion MRI data acquired for *K* = 200 healthy adults (age 21–36, 48.5% female) participating in the Human Connectome Project [30]. Structural brain networks were mapped at several spatial resolutions (*N* = 256, 360, 512 regions; *Materials and Methods, Connectome mapping*). For each individual, the resulting weighted adjacency matrix was thresholded at 10%, 15% and 20% connection density to eliminate potentially spurious connections [31], and subsequent analyses were carried out on the obtained weighted connectomes. Communication matrices quantifying the communication efficiency between every pair of regions were computed (Fig. 2a, *Materials and Methods, Network communication models*) and used to derive measures of send-receive asymmetry in communication efficiency (*Materials and Methods, Communication asymmetry test*). We focus on describing the results for *N* = 360, corresponding to a state-of-the-art cortical atlas derived from high-quality multi-modal data from the HCP [32], and 15% connection density. Results for other connection densities and parcellation resolutions can be found in the Supplementary Information.

Significant asymmetries in the efficiency of sending versus receiving information were evident for most cortical regions (Fig. 3a,d,g). Regional values of send-receive asymmetry were significantly correlated across regions among the communication measures investigated (navigation and diffusion: *r* = 0.29, navigation and search information: *r* = 0.31, diffusion and search information: *r* = 0.85; all *P* < 10^−7^). Based on these send-receive asymmetries, we classified all regions as senders, receivers or neutral. As expected from the strong correlation between them, diffusion and search information asymmetries led to similar classifications, likely due to their mutual dependence on random walk processes. While communication under navigation is guided by different mechanisms, classification consistency across measures was greater than expected by chance (Supplementary Note 1).

**FIG. 3.**
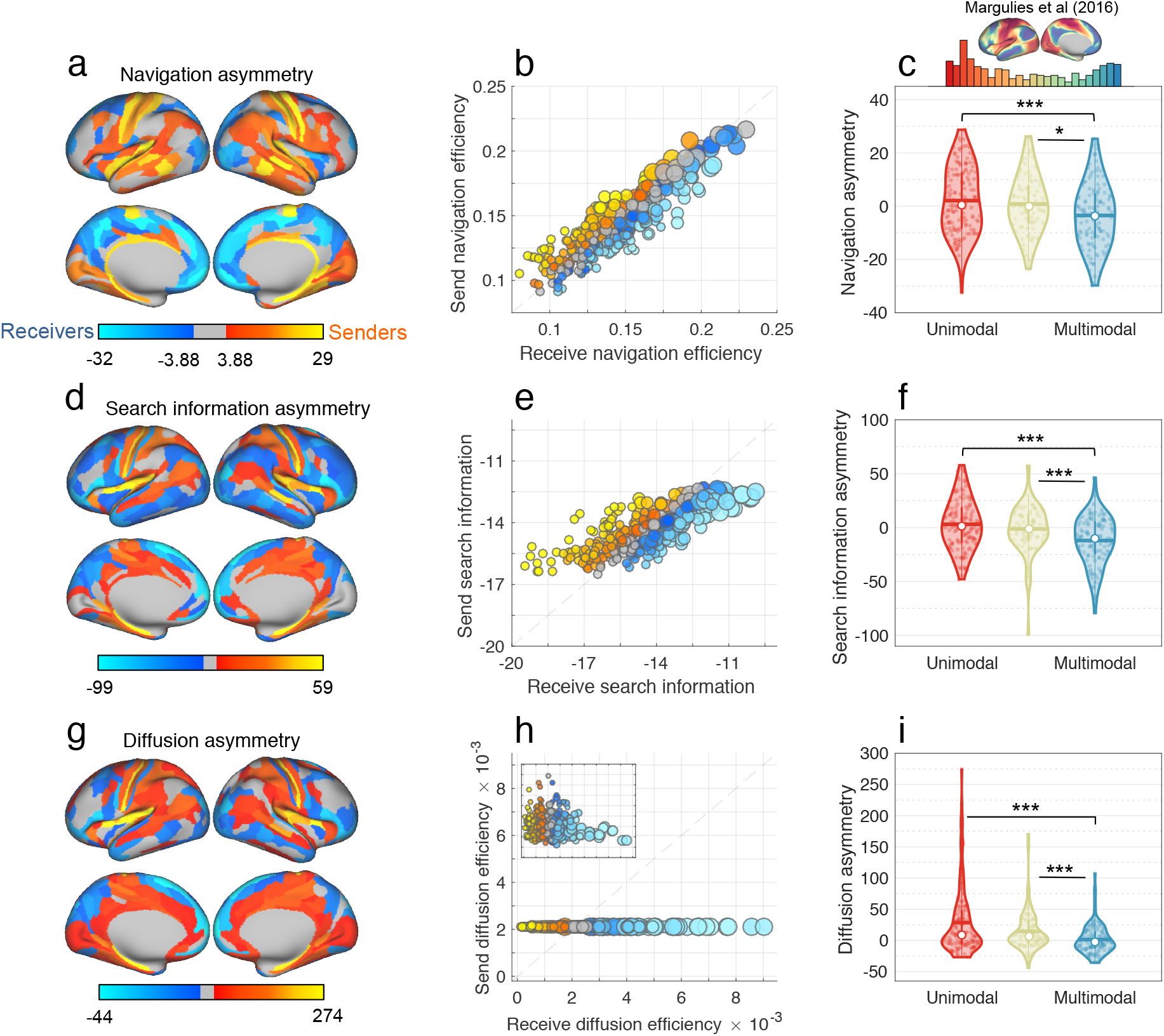
Send-receive communication asymmetry in the human connectome (*N* = 360 at 15% connection density). **(a)** Cortical projection of send-receive asymmetry values under navigation. Regions associated with significant send-receive asymmetry are classified as putative senders (orange) and receivers (blue). Regions colored gray are neutral and do not show significant send-receive asymmetry. **(b)** Scatter plot showing correlation between send (vertical axis) and receive (horizontal axis) efficiency across regions under navigation. Send and receive efficiency values were aggregated across all individuals for each region. Markers are colored according to send-receive asymmetry values (colors approximately match that of panel a). Small, medium and large markers represent nodes with low (*κ* < 50), medium (50 ≤ *κ* ≤ 100) and high (*κ* > 100) degree, respectively. The dashed line marks the *x* = *y* identity line. The distance between markers and the identity line provides a geometric interpretation of regional bias towards sending (*x* < *y*) or receiving (*x* > *y*) efficiencies. **(c)** Top: Distribution of the cortical gradient eigenvectors used as a measure of functional heterogeneity [35]. Bottom: Violin plots showing distribution of send-receive asymmetries for regions classified as unimodal (red), transitional, (beige) and multimodal (blue) regions. Horizontal bars and white circles denote, respectively, the mean and median of the distributions. Stars denote significant differences in between-group medians given by a two-sided Wilcoxon rank sum test (one, two and three stars denote *P* < 0.05,0.005,0.0005, respectively). **(d-e)** Search information equivalent of a-c. **(g-i)** Diffusion efficiency equivalent of a-c.

Primary sensory and motor regions were identified as senders (A1, S1 and M1 across all communication measures and V1 for navigation and diffusion). This is consistent with the notion that early auditory, visual and sensory-motor areas constitute the three main input streams to the cortex, being the first cortical regions to process sensory stimuli transmitted from subcortical structures [32, 33]. In contrast, expanses of the orbital and polar frontal cortices, the medial and dorsolateral prefrontal cortices, and the precuneus were classified as receivers. These regions have been proposed as putative functional hubs, supporting abstract and high-order cognitive processes by integrating multiple modalities of information [34–36]. Other regions consistently identified as senders included portions of the superior temporal, medial temporal and posterior cingulate cortices, while parts of intraparietal sulcus cortex, dorsal and ventral streams consistently ranked amongst receivers. The MT+ complex was also a prominent receiver, potentially reflecting the role of higher order sensory regions as integrators of signals transmitted from primary/earlier cortices. Certain regions were classified as senders under one communication measure but receivers under another measure, possibly reflecting how the three communication measures uniquely interact with connectome topology. Inconsistently classified regions included portions of the paracentral, cingulate, middle temporal and inferior temporal cortices. Details on how to access complete send-receive asymmetry tables and cortical maps are provided in *Materials and methods, Data availability*.

Despite significant asymmetries in the efficiency of sending versus receiving information *within* individual regions, these send-receive asymmetries were superposed atop a strong correlation across regions between send and receive efficiency (navigation: *r* = 0.95, search information: *r* = 0.79; both *P* < 10^−15^; Fig. 3b,e). In other words, efficient senders were also typically efficient receivers, and vice versa. Therefore, while all senders were by definition significantly more efficient at sending than receiving, some senders were in fact less efficient at sending than some receivers. In contrast, send and receive efficiencies were not correlated under diffusion (*r* = −0.1, *P* = 0.1). In addition, send efficiency was relatively uniform across regions under diffusion, while receive efficiency showed markedly higher regional diversity.

Node degree and participation (Supplementary Note 2) were associated with send-receive asymmetries under diffusion and search information (degree: *r* = −0.54, −0.70, participation: *r* = −0.29, −0.32, respectively. All *P* < 10^−7^), with senders and receivers characterized by low and high degree/participation, respectively. We regressed out the influence of degree on send-receive asymmetry and found that primary cortices remained senders while frontal and prefrontal regions remained receivers. We also noticed that sensory-motor and auditory cortices displayed a significantly higher propensity towards outgoing communication than expected based on their degree alone (Supplementary Fig. 2). Send-receive asymmetry under navigation was not correlated with degree or participation (both *P* > 0.05), with senders and receivers uniformly distributed across the degree distribution.

### Senders and receivers situated within cortical gradients

Next, we aimed to test whether regional variation in send-receive asymmetry would accord with established hierarchies of cortical organization [37]. To this end, we investigated whether senders and receivers would reside at opposing ends of a previously delineated uni-to multimodal gradient of functional connectivity [35]. Under all three communication measures, senders were more likely to be located at the unimodal end of the gradient, whereas the multimodal end was occupied by receivers. More specifically, send-receive asymmetry and the cortical gradient were significantly correlated across regions (*r* = −0.20, −0.30, −0.29, for navigation, diffusion and search information, respectively. All *P* < 10^−4^). These associations remained significant after accounting for the influence of degree on send-receive asymmetries (all *P* < 2 × 10^−4^).

In further analyses, regions were classified as unimodal (U), transitional (T) or multimodal (M) based only on the cortical gradient (*Materials and Methods, Cortical gradient of functional heterogeneity*). We compared the send-receive asymmetry of these groups and found that unimodal and transitional areas were significantly more efficient at outgoing communication compared to multimodal areas (Fig. 3c,f,i; *P*_*T*>*M*_ = 0.01, 2 × 10^−4^, 2 × 10^−4^ and *P*_*U*>*M*_ =4 × 10^−4^, 3 × 10^−6^, 8 × 10^−7^, for navigation, search information and diffusion, respectively). Send-receive asymmetry did not differ between unimodal and transitional regions.

These results were generally robust to variations in cortical parcellation and connection density thresholds (Supplementary Figs. 3 and 4). Taken together, our findings demonstrate that decentralized communication measures applied to the undirected human connectome unveil regional distinctions between putative senders and receivers. Furthermore, we show that knowledge about the direction of information flow can elucidate novel organizational structure within established cortical hierarchies, such as the biases towards outgoing and incoming communication efficiency of uni- and multimodal regions, respectively.

**FIG. 4.**
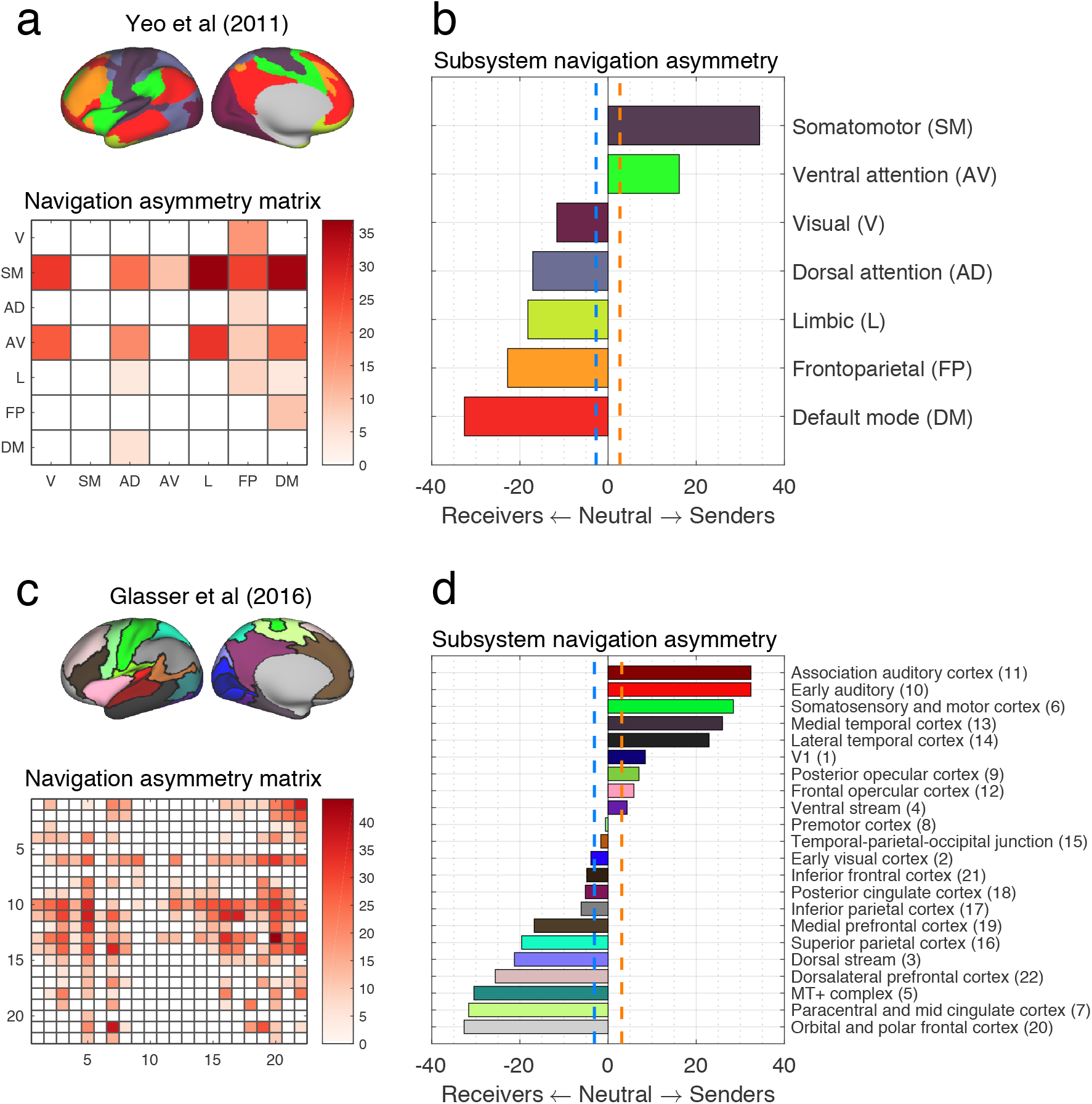
Send-receive asymmetry of cortical subsystems under navigation (*N* = 360 at 15% connection density). **(a)** Projection of *M* = 7 distributed resting-state networks onto the cortical surface (top) and send-receive asymmetry matrix under navigation (bottom). A matrix element *A*(*i,j*) > 0 denotes that communication occurs more efficiently from *i* to *j* than from *j* to *i*. Send-receive asymmetry values that did not survive multiple comparison correction were suppressed and appear as white cells. For ease of visualization and without loss of information (since *A*(*i,j*) = −*A*(*j, i*)), negative values were omitted. **(b)** Resting-state networks ranked by propensity to send (top) or receive (bottom) information. Dashed vertical lines indicate a significant bias towards outgoing (orange) and incoming (blue) communication efficiency. **(c-d)** Same as (a-b), but for *M* = 22 spatially contiguous cortical subsystems. Numbers listed next to module names identify corresponding rows and columns in the asymmetry matrix.

### Senders and receivers of cortical subsystems

Next, we sought to investigate pairwise send-receive asymmetries between large-scale cortical subsystems (Supplementary Fig. 5). We assigned cortical regions to distributed cognitive systems according to established resting-state networks comprising *M* = 7 and 17 subsystems [38]. In addition, we employed a multimodal partition of the cortex into *M* = 22 spatially contiguous subsystems [32]. Regional communication matrices were downsampled to subsystem resolution and send-receive asymmetries were computed at individual and pairwise subsystem levels (*Materials and Methods, Cortical sub-systems*).

In keeping with the regional findings, send-receive asymmetry matrix values were significantly correlated across subsystems among the communication measures investigated (e.g., navigation and diffusion: *r* = 0.60, navigation and search information: *r* = 0.66, diffusion and search information: *r* = 0.96; all *P* < 10^−15^, *M* = 17). Given the consistency of findings across communication measures, we focus on navigation in this section (Fig. 4). Complete results for navigation, diffusion and search information are shown, respectively, in Supplementary Figs. 6, 7 and 8.)

As shown in Fig. 4a,b, the somatomotor and ventral attention networks were the most prominent senders for the *M* = 7 partition. Prominent receivers included the default mode, frontoparietal and limbic networks, which were more efficiently navigated from a number of cognitive systems than vice versa. Interestingly, adopting a higher resolution functional partition (*M* = 17) suggested that sub-components of coarse (*M* = 7) resting-state networks can assume different roles as senders and receivers. For instance, the visual network was segregated into early (e.g., V1 and V2) and late areas of the visual cortex (e.g., MT+ complex and dorsal and ventral streams), with the first being a sender and the latter a receiver (Supplementary Fig. 6). Other systems that exhibited this behaviour included the ventral attention, limbic, somatomotor and default mode networks. These findings reiterate that, despite the presence of asymmetries in send-receive efficiency, cognitive systems are not exclusively capable of sending or receiving, suggesting connectome topology may allow for context-dependent directionality of neural information flow between functional networks.

We also identified senders and receivers for a high-resolution cortical partition comprising *M* = 22 subsystem [32]. This enabled a fine-grained, yet visually interpretable, characterization of send-receive asymmetries (Fig. 4c). Cortical domains associated with auditory, somatosensory and motor processes ranked amongst the strongest senders, while frontal and prefrontal areas consistently featured amongst the most prominent receivers (Supplementary Figs. 6 to 9). Together, these results provide putative multi-scale maps of how the structural substrate of the human connectome may facilitate directional information flow between cognitive subsystems.

### Senders, receivers and effective connectivity

We sought to validate our characterization of subsystems as senders or receivers using an independent data modality. To this end, time series summarizing the functional dynamics of cortical subsystems were extracted from resting-state functional MRI data for the same *K* = 200 HCP participants. For each individual, we used spectral DCM [39, 40] to compute effective connectivity between cortical subsystems (*M* = 7,17, 22, see *Materials and Methods, Effective connectivity*). Pairwise effective connectivity asymmetry was computed at the scale of subsystems by applying the previously described asymmetry test to the estimated effective connectivity matrices (Fig. 2c). Importantly, effective connectivity is an inherently directed (asymmetric) measure of connectivity. This allowed us to test whether send-receive asymmetries in communication efficiency (derived from diffusion MRI) and effective connectivity (derived from resting-state fMRI) are correlated (Fig. 2d).

Communication and effective connectivity send-receive asymmetries were significantly correlated across pairs of subsystems (Fig. 5). These associations were significant for all three communication measures and were replicated across two independent resting-state functional MRI sessions and multiple structural connection densities. For instance, for *M* = 17, fMRI session 1 and 15% connection density, we found *r* = 0.51,0.32,0.32 for navigation, diffusion and search information, respectively (all *P* < 10^−4^). Similarly, for *M* = 22, fMRI session 2 and 15% connection density, we obtained *r* = 0.45, 0.48, 0.48 for navigation, diffusion and search information, respectively (all *P* < 10^−12^). No significant correlations were found for *M* = 7, possibly due to the lack of statistical power afforded by only 21 data points comprising the upper triangle of asymmetry matrices. These results suggest that biases in the directionality of neural signalling inferred from the structural connectome are related to the directions of causal functional modulation during rest. Therefore, they establish a correspondence between structural (connectome topology and network communication measures) and functional (effective) directions of neural information flow.

**FIG. 5.**
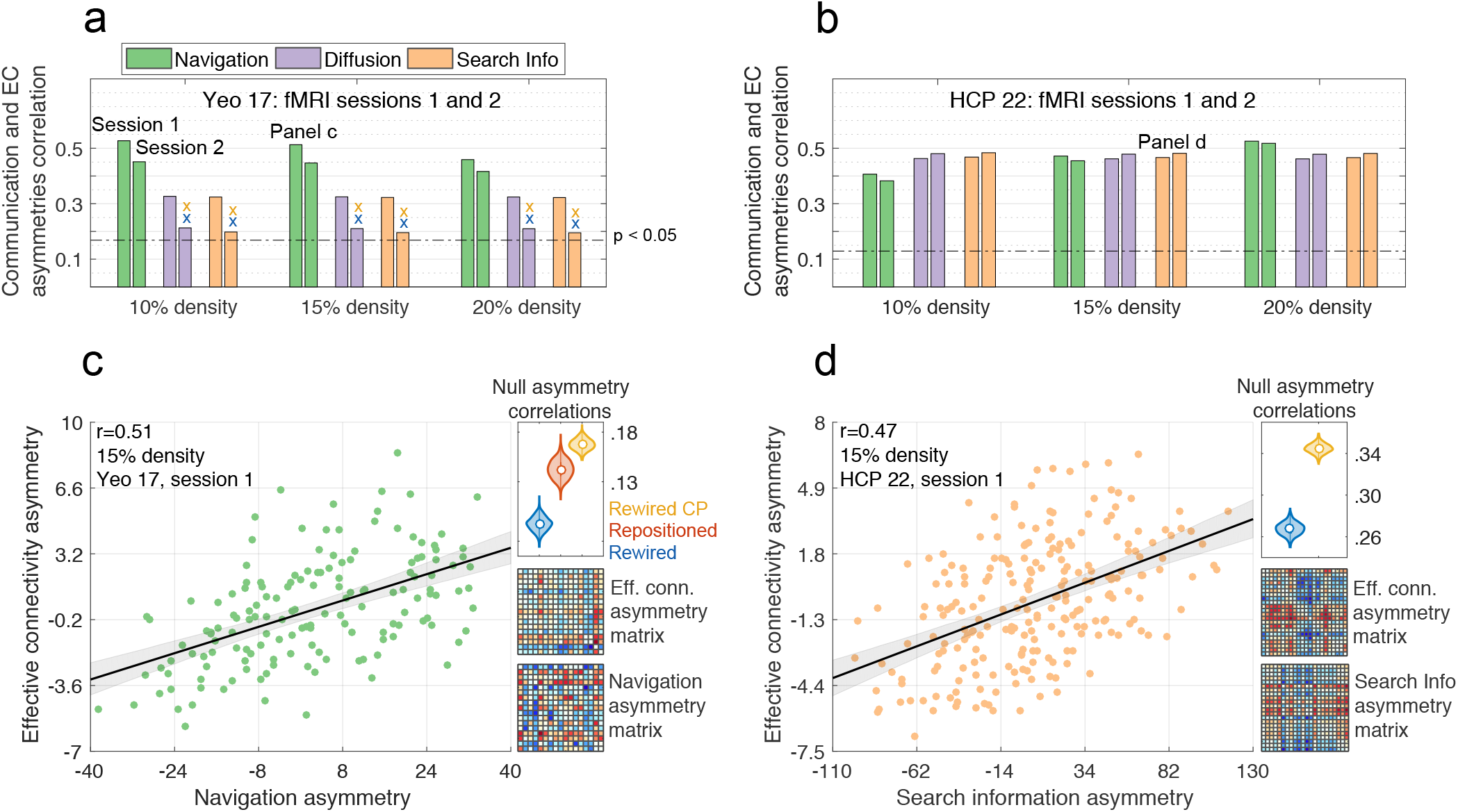
Relationship between send-receive asymmetry and directionality of effective connectivity (*N* = 360). **(a)** Bars denote the Pearson correlation coefficients between send-receive and effective connectivity asymmetries for *M* = 17 cortical subsystems. Bars are colored according to the three communication measures: i) navigation (green), ii) diffusion (violet), and iii) search information (beige). Correlations were computed for two independent resting-state fMRI sessions (Sessions 1 and 2) and multiple structural connection density thresholds (10, 15 and 20%). Significance threshold of *P* < 0.05 is indicated with a dotted line. Crosses mark associations that were *not* statistically stronger than those found in families of 1000 rewired (blue) and 1000 cost-preserving rewired (yellow) connectomes, respectively (repositioned connectomes, relevant for navigation, led to statistically weaker associations in all scenarios). **(b)** Replication of Panel **a** for a cortical partition comprising *M* = 22 subsystems. **(c)** Left: scatter plot illustrating the correlation between navigation and effective connectivity asymmetries for *M* = 17, 15% connection density and fMRI session 1. Shadows denote the 95% bootstrapped confidence interval. Top-right: Distribution of correlations obtained for families of 1000 rewired (blue), repositioned (red) and cost-preserving rewired (yellow) connectomes. Bottom-right: Send-receive asymmetry matrices for effective connectivity (resting-state functional MRI) and navigation (diffusion MRI). The upper-triangular elements of these two matrices were correlated to test whether senders and receivers were consistently identified across independent modalities. **(d)** Replication of Panel **c** for search information, *M* = 22, 10%) connection density and fMRI session 2. EC: effective connectivity; *r*: Pearson correlation coefficient.

We sought to determine whether the above association between communication and effective connectivity could be explained by certain properties of connectome organization. We generated ensembles of randomized connectomes in which (i) connectome topology was rewired while preserving degree distribution [41]; (ii) connectome topology was rewired while preserving degree distribution and total network cost (defined as the sum of Euclidean distances between structurally connected nodes [14]); and (iii) nodes were spatially repositioned while preserving topology (relevant only for navigation; see Supplementary Note 3). For all families of randomized connectomes, correlations between asymmetries in effective connectivity and communication efficiency were significantly decreased compared to empirical results (e.g., Fig. 5c,d top-right comer, all *P* < 10^−3^; with the exception of diffusion and search information for the *M* = 17 partition in fMRI session 2, Fig. 5a). These results indicate that the relationship between send-receive asymmetry and directionality of effective connectivity cannot be explained by the combination of connectome degree distribution and network cost, since these properties were preserved in random ensembles (i) and (ii). For navigation, random ensemble (iii) highlights the importance of connectome geometry in addition to topology.

### Control analyses

Having established several properties of the senders and receivers of the human connectome, we aimed to determine whether our results were robust to alternative definitions of send-receive communication asymmetry and changes in our connectome mapping pipeline. First, we redefined our communication asymmetry measure using non-parametric Wilcoxon rank sum tests instead of t-tests (Supplementary Note 4). This approach ensures that send-receive asymmetries are robust to deviations from normality and outliers. Second, we computed the send-receive asymmetries of connectomes derived with probabilistic tractography (Supplementary Note 5). Third, we investigated send-receive asymmetries in connectomes including subcortical structures (Supplementary Note 6). Send-receive asymmetries were compatible across all three scenarios and remained consistently associated with the directionality of effective connectivity (Supplementary Fig. 10). Interestingly, while the classification of nodes into senders and receivers in connectomes containing subcortical structures remained largely unaltered, biases towards incoming or outgoing communication were less pronounced (Supplementary Figs. 12 and 13), suggesting a role of subcortical structures as mediators of neural signalling directionality.

### Senders and receivers of non-human connectomes

The association between effective connectivity directionality and the send-receive asymmetry of undirected connectomes indicates that signalling directions in the human brain are not exclusively determined by axonal directions. To further quantify this observation, we next sought to establish the extent to which signalling directionality is determined by axonal directions *per se*, compared to other potential determining factors such as network topology and geometry.

Invasive connectome reconstruction techniques allow for the resolution of axonal directionality, producing directed connectomes for a host of non-human species [42]. Here we consider the connectomes of the fruit fly (*drosophila*) [43, 44], mouse [45, 46] and macaque [47] (*Materials and methods, Non-human connectomes*). We began by computing the send-receive asymmetries of these directed connectomes. In this case, communication asymmetry is introduced both by the asymmetric character of the network communication measures and by the presence of directed connections. Next, we symmetrized the connectomes by removing connection directionality, so that all connections could be traversed bidirectionally (*Materials and methods, Symmetrized non-human connectomes*), and recomputed send-receive asymmetries for the resulting undirected networks. In this scenario, as with human undirected connectomes, asymmetries are introduced solely by the asymmetry inherent to the network communication measures. We tested whether send-receive asymmetry values computed in the directed (original) and undirected (symmetrized) non-human connectomes were correlated across regions. Evidence of a correlation would suggest that the undirected topology and geometry of connectomes are influential in determining the directionality of neural signalling in the absence of directed connections.

We found that undirected and directed send-receive asymmetries were correlated for binarized (fly: *r* = 0.95,0.96, mouse: *r* = 0.58,0.50, macaque: *r* = 0.87, 0.75, for diffusion and search information asymmetries, respectively; Fig. 6a,c,d) and weighted (fly: *r* = 0.58,0.84, 0.41, mouse: *r* = 0.34, 0.32, 0.38, macaque: *r* = 0.67,0.80, −0.26, for navigation, diffusion and search information asymmetries, respectively; Fig. 6b,e,f) nonhuman connectomes. All reported *r* had *P* < 10^−10^ and thus survived Bonferroni correction for multiple comparisons. The exception was the association for the macaque weighted search information (*P* = 0.01), potentially indicating the spurious nature of this negative correlation. It is worth noting that binary navigation paths are seldom asymmetric for densely connected networks such as nonhuman connectomes, resulting in weak/undefined correlations between directed and undirected send-receive asymmetries. In addition, due to the presence of connection weight asymmetries between bidirectionally connected node pairs, original and symmetrized connectomes are more similar for binary than weighted networks, explaining the stronger associations observed for binarized connectomes.

**FIG. 6.**
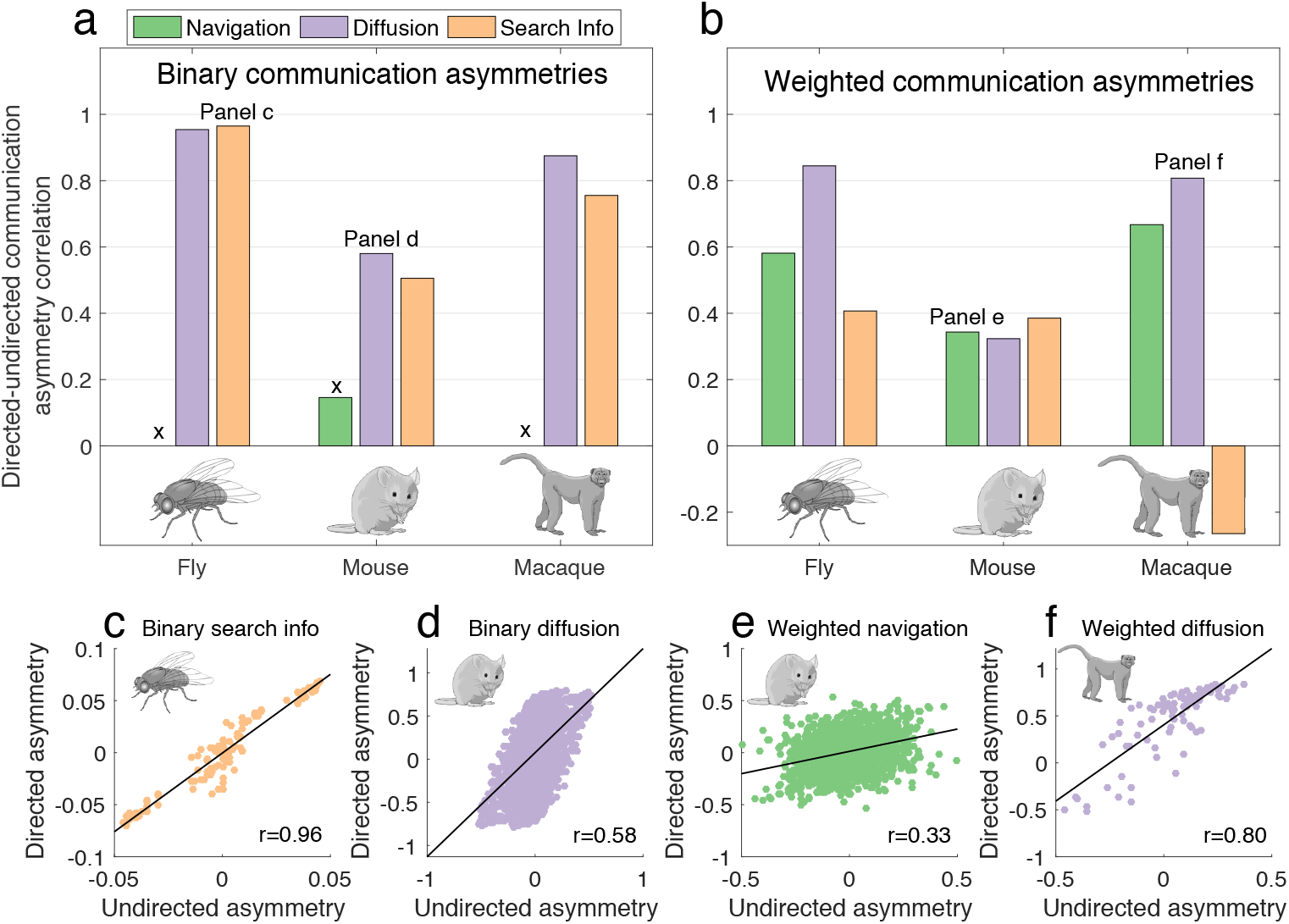
Directed and undirected (symmetrized) send-receive asymmetries in non-human connectomes. **(a)** Pearson correlation coefficient across regions in send-receive asymmetry values between directed and undirected navigation (green), diffusion (violet) and search information (beige) asymmetries for binarized (unweighted) connectomes. Black crosses indicate *nonsignificant* (*P* > 0.05) or undefined (in the case of lack of communication asymmetry) correlations. **(b)** Same as (a), but for weighted connectomes. **(c-f)** Scatter plots illustrating the association between directed and undirected communication asymmetries (*r*: Pearson correlation coefficient, *P*: associated P-value).

Finally, we observed that regional senders and receivers of undirected (symmetrized) non-human connectomes also recapitulated putative hierarchies of functional specialization (Supplementary Note 7). For instance, macaque sensory, visual and motor areas were senders, while portions of the frontal and prefrontal cortices were receivers (Supplementary Figs. 14g,h,i and 15). Collectively, these findings provide further evidence that the directionality of neural signalling is partially determined by the undirected architecture of nervous systems across species.

## DISCUSSION

The present study focused on characterizing the directionality of neural information flow arising from the application of decentralized network communication measures to connectomes. In a recent study, Avena-Koenigsberger and colleagues presented a first account of differences between send and receive communication in brain networks [16]. Here, we build on these efforts by contributing a statistical framework to compute send-receive communication asymmetry. We apply this framework to identify putative sender and receiver brain regions, as well as pairwise maps of neural signalling directionality for the nervous systems of several species.

Send-receive asymmetry recapitulated hierarchical patterns of cortical organization from a structural connectivity standpoint. Several studies of axonal tract-tracing and non-human connectomes [34, 48, 49], macroscale gradients of cortical organization [35, 37, 50], and computational models of neuronal dynamics [27, 51–54] converge to a common conceptualization of a cortical hierarchy of functional specialization. The bottom of the hierarchy tends to comprise high-frequency, low-degree, unimodal, sensory and motor areas that constitute the main inputs of perceptual information to the brain. At the top, low-frequency, high-degree, multimodal regions are conjectured to integrate multiple streams of information in order to support higher cognitive functions. Our observations of a send-receive spectrum of cortical regions and subsystems complements this description of neural organization, placing senders and receivers, respectively, at the uni- and multimodal ends of the hierarchy.

Previous studies have demonstrated that navigation efficiency, search information and diffusion computed on structural connectomes are capable of inferring resting-state functional connectivity [8, 15, 24]. Here, we provided further evidence for the utility of decentralized communication models by showing an association between send-receive asymmetry—inferred from connectomes mapped with tractography and diffusion MRI— and directionality of effective connectivity—computed from spectral DCM applied to resting-state fMRI. This relationship was robust to variations in tractography algorithms, cortical subsystem parcellations, treatment of subcortical structures, send-receive asymmetry statistical tests, structural connection density thresholds and two independent resting-state fMRI sessions. Compared to the first fMRI session, the strength of this relationship was weaker in the second session for the case of the *M* = 17 cortical subsystems. This may be due to the effect of MRI phase-encoding differences between the two sessions (*Materials and Methods, Send-receive effective connectivity asymmetry*) on particular subsystem parcellations, although this requires further investigation.

Recent work has demonstrated the validity of spectral DCM in multi-site longitudinal settings [55] and using optogenetics combined with functional MRI in mice [56]. The use of spectral DCM instead of the traditional task-based DCM was motivated by two important factors. First, spectral DCM infers effective connectivity from resting-state fMRI data, allowing validation of our findings independent of hypotheses about the directionality of causal connectivity specific to certain task scenarios. In addition, recent evidence indicates that functional connectivity topology at rest shapes task-evoked fluctuations, highlighting the cognitive relevance of resting-state neural dynamics [57, 58]. Second, spectral DCM is capable of handling relatively large networks comprising many regions [59]. This enabled a direct comparison between asymmetries in send-receive efficiency and effective connectivity at the level of subsystems spanning the whole cerebral cortex. Our results provide cross-modal evidence that network communication measures accurately capture aspects of directional causal influences between neural systems. Structurally-derived communication asymmetry may help formulate hypotheses for DCM studies, potentially reducing the search space of candidate network models [60]. In addition, send-receive asymmetry may be useful in understanding asymmetric responses in functional dynamics following exogenous stimulation of brain regions [27, 52].

The analyses of human undirected connectomes indicate that meaningful patterns of neural signalling directionality can be inferred without knowledge of the directions of axonal projections. We provided further evidence for this notion by examining non-human connectomes, for which information on axonal directionality is invasively derived. Send-receive asymmetries computed for directed connectomes were significantly associated to those derived from networks for which the directionality of connections was suppressed. Moreover, senders and receivers computed from undirected version of non-human connectomes also recapitulated putative functional roles of brain regions. These results indicate that despite the documented importance of directed connections [47, 61], the undirected architecture of nervous systems also imposes constraints on signalling directionality. This may suggest the presence of fundamental, cross-species organization properties of brain networks that facilitate decentralized communication between neural elements. It is worth noting that send-receive asymmetry is more pronounced between pairs of regions that are not directly connected, for which communication takes place along multi-hop paths. Consequently, as formulated here, send-receive asymmetry is not well suited to perform inference for structurally connected nodes, and thus should not be conceptualized as a methodology to transform an undirected structural connectome into a directed graph. Future work exploring alternative formulations of communication asymmetry could attempt to infer directed structural traits from undirected connectomes.

Send-receive asymmetry is a result of the interaction between asymmetric network communication models and the topology and geometry of brain networks. Navigation depends on local knowledge of the network’s spatial embedding to identify communication paths, while diffusive processes rely solely on local connectivity knowledge to propagate signals. Despite these conceptual differences, navigation efficiency, search information and diffusion efficiency led to similar patterns of send-receive asymmetry. The classification of cortical regions into senders and receivers, as well as the association with effective connectivity directionality was generally consistent across measures. This indicates that our results may be primarily driven by how the architecture of brain networks gives rise to general patterns of communication asymmetry, rather than by specific strategies of neural signalling. An interesting deviation from these consistencies was observed in the relationship between regional send and receive efficiencies. While navigation and search information showed a positive correlation between send and receive efficiencies, this was not the case for diffusion. Moreover, as previously reported [16], diffusion receive efficiency showed markedly greater regional variation compared to diffusion send efficiency. This is a consequence of high degree nodes being more accessible to incoming random walkers than low degree ones.

We reiterate that a significant send-receive asymmetry does not preclude information transfer in a particular direction, in the same way that regions classified as senders (receivers) are capable of receiving (sending) information. Interestingly, we also found that coarse functional networks with significant biases towards incoming or outgoing communication are typically comprised of subcomponents placed along different positions of the sender-receive spectrum. This may facilitate feedback loops in which high-order regions send information to sensory cortices, allowing for flexible and context-dependent transfer of neural information. These results support the notion that cortical computations do not follow a strictly serial paradigm, but rather involve distributed hierarchies of parallel information processing [38, 48].

Several limitations of the present study should be considered. Send-receive asymmetry was defined statistically across subjects. Future developments are necessary to conceptualize robust measures of subject-level send-receive asymmetry. In addition, alternative asymmetric network communication measures such as Markovian queuing networks [62], linear transmission models of spreading dynamics [19, 20] and cooperative learning [63] can lead to further insight into the large-scale directionality of neural signalling. Additional measures of directed functional connectivity such as transfer entropy and Granger causality may offer supplementary cross-modal validation of send-receive asymmetry. Importantly, tractography algorithms are prone to known biases, potentially influencing results regarding human structural connectomes [31, 64, 65]. Lastly, navigation was computed based on the Euclidean distance between brain regions. Alternative distance measures taking it account axonal fiber length may provide more biologically realistic guidance for connectome navigation.

In conclusion, we showed that the large-scale directionality of neural signalling can be inferred, to a significant extent, from the interaction between decentralized network communication measures and the undirected topology and geometry of brain networks. These results challenge the belief that connectomes mapped from *in vivo* diffusion data are unable to characterize asymmetric interactions between cortical elements. Our findings introduce decentralized network communication models as a new avenue to explore directional functional dynamics in human and non-human connectomes.

## MATERIALS AND METHODS

### Connectivity data

Minimally preprocessed diffusion-weighted MRI data from 200 healthy adults (age 21–36, 48.5% female) was obtained from the Human Connectome Project (HCP) [30]. Details about the acquisition and preprocessing of this diffusion MRI are described in [66, 67].

Connectome analyses are sensitive to the number of nodes used to reconstruct brain networks [68]. We aimed to reproduce our key findings for human connectomes constructed with different granularities of cortical segmentation comprising *N* = 256, 360, 512 regions/nodes. The parcellations for *N* = 256, 512 segment the cortex into approximately evenly sized regions that respect predefined anatomical boundaries. Details on the construction of these parcellations are described in [15]. In addition, we mapped connectomes using the HCP MMP1.0 atlas (*N* = 360), a cortical parcellation based on multimodal data from the HCP [32].

Diffusion tensor imaging combined with a deterministic tractography pipeline was used to map connectomes for each individual. Deterministic tractography leads to less false positive connections than other reconstruction methods, and thus may better suit connectome mapping compared to alternative tractography methods [31, 64, 65]. Computations were carried out using MR-trix3 [69] with the following parameters: FACT tracking algorithm, 5 × 10^6^ streamlines, 0.5 mm tracking step-size, 400 mm maximum streamline length and 0.1 FA cutoff for termination of tracks. Connection strength between a pair of regions was determined as the number of streamlines with extremities located in the regions divided by the product of the surface area of the region pair, resulting in a *N* × *N* weighted connectivity matrix per subject. For each individual, the resulting weighted adjacency matrix was thresholded at 10%, 15% and 20% connection density to eliminate potentially spurious connections [31], and subsequent analyses were carried out on the obtained weighted connectomes.

The fruit fly connectome was mapped using images of 12,995 projection neurons in the female *drosophila* brain available in the FlyCircuit database [43, 44]. Single neurons were labeled with green fluorescent protein and traced from whole brain three-dimensional images. Individual neurons were grouped into 49 local processing units with specific morphology and function. The resulting connectome is a 49 × 49 weighted, directed, whole-brain network for the fruit fly, with 83% connection density.

The Allen Institute for Brain Science mapped the mesoscale topology of the mouse nervous system by means of anterograde axonal injections of a viral tracer [45]. Using two-photon tomography, they identified axonal projections from the 469 injections sites to 295 target regions. Building on these efforts, Rubinov and colleagues constructed a directed, bilaterally symmetric, whole-brain network for the mouse, comprising *N* =112 cortical and subcortical regions with 53% connection density [46]. Connections represent interregional axonal projections and their weights were determined as the proportion of tracer density found in target and injected regions.

Markov and colleagues applied 1615 retrograde tracer injections to 29 of the 91 areas of the macaque cerebral cortex, spanning occipital, temporal, parietal, frontal, prefrontal and limbic regions [47, 70]. This resulted in a 29 × 29 weighted, directed, interregional sub-network of the macaque cortico-cortical connections with 66% connection density. Connection weights were estimated based on the number of neurons labelled by the tracer found in source and target regions, relative to the amount found in whole brain.

### Network communication measures

A weighted connectome can be expressed as a matrix *W* ∈ ℝ^*N × N*^, where *W_ij_* is the connection weight between nodes *i* and *j*. Connection weights are a measure of similarity or affinity, denoting the strength of the relationship between two nodes (e.g., streamline counts in tractography or fraction of labelled neurons in tract tracing). The computation of communication path lengths mandates a remapping of connection weights into lengths, where connection lengths are a measure of the signalling cost between two nodes [11]. The transformation *L* = −log_10_(*W*/(*max*(*W*) + *min*(*W*_>0_)) ensures a monotonic weight-to-length remapping that attenuates extreme weights [8, 71], where *min*(*W*_>0_) denotes the smallest positive value in *W*, preventing the remapping of the maximum value of *W* to 0.

Navigation (also referred to as greedy routing) is a decentralized network communication model that utilizes information about the network’s spatial embedding to route signals without global knowledge of network topology [28]. Navigation is reported to achieve near-optimal communication efficiency in a range of real-world complex networks, including the connectomes of several species [15, 17, 18].

Navigation from node *i* to *j* was implemented as follows. Progress to *i*’s neighbour that is closest in distance to *j*. Repeat this process for each new node until *j* is reached—constituting a successful navigation path—or a node is revisited—constituting a failed navigation path. The distance between two nodes was computed as the Euclidean distance between the centroids of their respective gray matter regions. For each parcellation resolution, a single Euclidean distance matrix was computed in standard space and utilized to guide the navigation of each individual’s connectome.

Let Λ denote the matrix of navigation path lengths. If node *i* cannot navigate to node *j*, Λ_*ij*_ = ∞. Otherwise, Λ_*ij*_ = *L_iu_* +… + *L_vj_*, where {*u*,…, *v*} is the sequence of nodes visited during navigation. Note that while navigation paths are identified based on the Euclidean distance between nodes, navigation path lengths are computed in terms of connection lengths derived from the structural connectivity matrix *W*. Navigation efficiency is given by *E_nav_*(*i, j*) = 1/Λ_*ij*_, where *E_nav_*(*i, j*) is the efficiency of the navigation path from node *i* to *j* [15].

A diffusion process is a network communication model whereby information is broadcast along multiple paths simultaneously [22]. Diffusion can be understood in terms of agents, often termed random walkers, which are initiated from a given region and traverse the network independent of each other by randomly selecting a connection to follow out from each successive region that is visited. Diffusive communication does not mandate assumptions on global knowledge of network topology, constituting, from this perspective, a biologically plausible model for neural communication [9]. Diffusion efficiency [23] is related to how many intermediate regions (synapses), on average, a naive random walker needs to traverse to reach a desired destination region.

Let *T* denote the transition probability matrix of a Markov chain process with states corresponding to nodes in the adjacency matrix *W*. The probability of a random walker at node *i* stepping to node *j* is given by 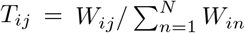. The expected number of hops 〈*H_ij_*〉 a random walker takes to travel from node *i* to node *j* is given by [72]:

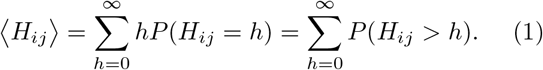

This result is given by the fact that the expected value of a random variable is given by the sum of its complementary cumulative distribution. The probability of a walker requiring more than *h* hops to reach node *j* is equal to the sum of the probabilities of the walker being at any node other than *j* after exactly *j* hops. To compute this, we define *T_j_* as the matrix *T* with all elements in the *j*th column set to zero, so that it is impossible for a walker to arrive at node *j*. This way, we have 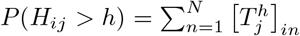 where 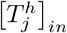 expresses the probabilities of walkers departing from *i* and reaching any other node expect *j* in exactly *h* hops. It follows that

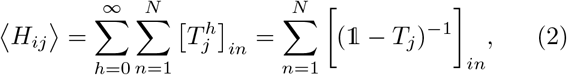

with the last derivation step following from the summation of an infinite geometric sequence. Further details on this derivation can be found in Refs. [11, 23, 72]. The diffusion efficiency communication matrix is given by *E_dif_*(*i, j*) = 1/*H_ij_*, where *E_dif_*(*i, j*) quantifies the efficiency of information flow from node *i* to node *j* under a diffusive process [23].

Search information relates to the probability that a random walker will serendipitously travel between two nodes via their shortest path [29], quantifying the extent to which efficient routes are hidden in the network topology. Previous studies suggest node pairs with an accessible shortest path—characterized by low search information—tend to show stronger resting-state functional connectivity [8].

The connection length matrix *L* can be used to compute Ω, where Ω_*ij*_ = {*u,…, v*} denotes the sequence of nodes traversed along the shortest path from node *i* to node j. The search information from *i* to *j* is given by *SI_ij_* = −*log_2_*(*P*(Ω_*ij*_)), where *P*(Ω_*ij*_) = *T_iu_* +… + *T_vj_* and *T* is the transition probability matrix. We define communication efficiency under search information as *E_si_*(*i, j*) = −*SI_i,j_*. This way, *E_si_*(*i, j*) quantifies the accessibility of the Ω_*ij*_ shortest path under diffusive communication.

### Send-receive communication asymmetry measures

Send-receive communication asymmetry matrices *A* ∈ ℝ^*N* × *N*^ were computed as detailed in *Measures of send-receive communication asymmetry*. For each pair of regions or subsystems, a one-sample t-test was used to assess whether the mean of the send-receive asymmetry values across all individuals was significantly different from 0 (see *Supplementary Note 3* for an investigation of normality assumptions involved in t-tests and send-receive asymmetries based on non-parametric statistics). Bonferroni correction was then performed to control for the *N*(*N* − 1) /2 multiple comparisons corresponding to distinct pairs of regions. This was repeated for each of the three communication measures.

The communication asymmetry matrix *A* refers to pairwise asymmetric interactions between regions. We performed a similar test to derive a regional (i.e., nodewise) measure of send-receive asymmetry. Let {*S, R*} ∈ ℝ^*N* × *K*^ denote, respectively, the average send and receive efficiencies of nodes in the network such that 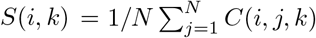 and 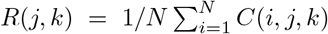. The difference between outgoing and incoming communication efficiencies of node *i* is given by *δ*(*i,k*) = *S*(*i,k*) − *R*(*i,k*). Analogous to the pairwise asymmetry test, we performed a one-sample t-test to determine whether the mean of the distribution *δ*(*i,k* = 1…*K*) is significantly different to 0. The resultant t-statistic, termed *a*(*i*), quantifies the communication asymmetry of node *i* by taking into account all of its incoming and outgoing communication efficiencies. Nodes with significant and positive (negative) *a* were classified as senders (receivers), while non-significant values of a were characterized neutral nodes. For each network communication measure, Bonferroni correction was performed to control for multiple comparisons across the *N* regions.

Regionally-aggregated send and receive efficiencies depicted in the scatter plots of Fig. 3e,h were computed as 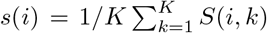 and 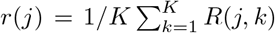, respectively. For navigation (Fig. 3b), we display the median send and receive efficiencies in order to attenuate outlier efficiency values and aid visualization.

Non-human directed connectomes were constructed from the results of numerous invasive experiments, often combining experiments across multiple animals of a given species to yield a single, representative connectome. As a result, non-human brain networks were not available for multiple individuals, precluding use of the communication asymmetry test defined for human connectomes. As an alternative, for non-human brain networks, we computed the communication asymmetry between nodes *i* and *j* as *A*(*i, j*) = (*E*(*i, j*) − *E*(*j, i*))/(*E*(*i, j*) + *E*(*j, i*)), where *E* is a communication efficiency matrix. While this measure does not constitute a statistical test of communication asymmetry, it allows us to evaluate differences in the directionality of information flow of non-human nervous systems.

### Cortical gradient of functional heterogeneity

Margulies and colleagues applied a diffusion embedding algorithm to resting-state fMRI data to identify latent components describing maximum variance in cortical functional connectivity [35]. The obtained components, termed “gradients”, are conjectured to describe macroscale principles of cortical organization [37]. In particular, the resultant principal gradient (*G*_1_) separated unifrom multimodal regions, spanning a spectrum from primary sensory-motor areas on one end, to the regions comprising the default-mode network on the other. We used this gradient as a quantitative measure of cortical functional heterogeneity and compared it to regional send-receive communication asymmetries. To this end, we downsampled the gradient from vertex to regional resolution by averaging the values comprising each of the *N* = 256, 360, 512 cortical areas defined by the parcellations that we used. Regions were grouped into the unimodal (*G*_1_ ≤ −2), transitional (−2 < *G*_1_ < 2) and multimodal (*G*_1_ ≥ 2) groups shown in Fig. 3.

### Cortical subsystems

Yeo and colleagues proposed a widely-used partition of the cortical surface into 7 and 17 resting-state functional networks [38]. These networks constitute distributed (i.e., non-contiguous) functional communities that have been implicated in a wide range of cognitive demands, as well as in rest. Glasser and colleagues used multimodal HCP data to identify 360 cortical regions. Subsequently, they grouped these regions into 22 contiguous subsystems based on geographic proximity and functional similarities [32]. We use these definitions of cortical partitions to investigate send-receive communication asymmetry at the level of subsystems.

First, we transformed the Yeo partitions (*M* = 7,17) from vertex to regional resolution. This was achieved by assigning each of *N* = 360 cortical regions to the resting-state network with the largest vertex count within the vertices comprising the region. The HCP partition (*M* = 22) does not necessitate this step, since it is already defined in terms of the *N* = 360 of the Glasser atlas.

Second, we downsampled individual communication efficiency matrices from regional (*N* = 360) to subsystem resolution (*M* = 7,17, 22) by averaging the pairwise efficiency of nodes assigned to the same subsystem. For two subsystems *u* and *v*, we have

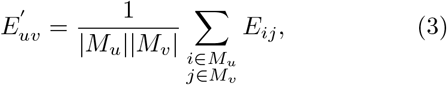

where *M_u_* and |*M_u_*| denote, respectively, the set and number of regions belonging to subsystem *u, E* ∈ ℝ^*N*×*N*^, and *E*′ ∈ ℝ^*M*×*M*^. Across *K* subjects, this results in a set of communication matrices *C* ∈ ℝ^*M* × *M* × *K*^ that is used to compute between-subsystems send-receive communication asymmetries as described in Fig. 2 and *Materials and Methods, Communication asymmetry test*.

The send-receive communication asymmetry for individual cortical subsystems was computed analogous to regional communication asymmetries as described in *Send-receive communication asymmetry measures*. For each network communication measure, Bonferroni correction for *M* and *M*(*M* − 1)/2 multiple comparisons was applied to individual and pairwise subsystems asymmetries, respectively.

### Send-receive effective connectivity asymmetry

Spectral DCM estimates effective connectivity from resting-state fMRI data. It receives as input time series characterizing the functional dynamics of neural activity and a network model describing how these elements are coupled. As opposed to the more common task-based DCM, spectral DCM estimates effective connectivity in the absence of experimental or exogenous inputs, characterizing functional modulations between neural elements based on intrinsic neural fluctuations at rest. Details on the generative models inherent to spectral DCM as well as the frequency-domain model inversion are described in [39, 59].

Minimally preprocessed resting-state fMRI data for the same *K* = 200 subjects was acquired from the HCP. Functional volumes were acquired during 14m33s at 720 TR, resulting in 1,200 time points. Data from two separate sessions (*rfMRI REST1 LR* and *rfMRI REST2 RL*, i.e., right-to-left and left-to-right encoding, performed on different days) was used to compute two estimates of effective connectivity for each subject. HCP acquisition and preprocessing of resting-state fMRI are detailed in [66, 67].

We computed the blood-oxygenation-level-dependent (BOLD) signal of *N* = 360 regions by averaging the time series of all cortical surface vertices comprised into a region. Next, the *N* regions were partitioned into *M* cortical subsystems as described in *Materials and Methods, Cortical subsystems*. For each subsystem, we performed a principal component analysis on all the time series belonging to it. The resultant first principal component was used to summarize the functional activity of a subsystem in a single time series. The *M* × 1, 200 time series of principal components were used as input to spectral DCM, together with a fully connected model of coupling strengths 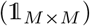, enabling estimation of effective connectivity between subsystems covering the whole cortex [59]. Spectral DCM estimations were carried out using SPM12.

Spectral DCM estimates signed effective connectivity, with positive and negative values indicating excitatory and inhibitory influences, respectively. Under the assumption that both excitatory and inhibitory processes are facilitated by communication between neural elements, we considered the absolute value of the estimated coupling strengths.

The obtained coupling strengths of each subject were concatenated. For each resting-state session, this yielded a *M* × *M* × *K* effective connectivity matrix, which were used to compute effective connectivity asymmetry between cortical subsystems, as described in Fig. 2 and *Materials and Methods, Communication asymmetry test*.

### Symmetrized non-human connectomes

Directed non-human connectomes (*W_d_*) were symmetrized in order to omit information on axonal directionality. Undirected (symmetric) networks (*W_u_*) were computed as 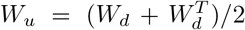, ensuring that all original connections in Wd can be traversed bidirectionally in *W_u_*. Directly connected node pairs do not show send-receive asymmetry under navigation, since both directions of routing will necessarily occur via the single connection linking the two nodes. For this reason, we restricted the analyses in *Senders and receivers in nonhuman connectomes* to node pairs that did not share a direct structural connection in *W_u_*. Non-human binary connectomes were constructed by discarding information on connectivity weight and considering only the presence or absence of directed connections. Formally, *B*(*i, j*) = 1 if *W*(*i, j*) ≠ 0 and *B*(*i, j*) =0 otherwise.

## Data availability

All analyses in the present study were carried out on publicly available datasets. Structural and effective human brain networks were mapped from Human Connectome Project data [30] (https://db.humanconnectome.org/). The fruit fly connectome was collated from data available in http://www.flycircuit.tw and can be found in the supplementary information of reference [44]. The macaque connectome was derived from data available at http://core-nets.org/ [47]. The mouse connectome was constructed from resources provided by the Allen Institute for Brain Science (https://mouse.brain-map.org/ [45]) and is available in the supplementary information of reference [46]. The cortical gradient of functional connectivity from reference [35] is available at https://www.neuroconnlab.org/data/index.html.

Send-receive communication asymmetry measures and other data necessary to generate key figures in this work will be made available through the BALSA database (https://balsa.wustl.edu/) upon manuscript acceptance.

## Code availability

Functions to compute navigation efficiency, diffusion efficiency and search information are available as part of the Brain Connectivity Toolbox (https://sites.google.com/site/bctnet/). Further analyses and computations were performed using MRtrix3 (www.mrtrix.org/), SPM12 (https://www.fil.ion.ucl.ac.uk/spm/software/spm12/) or custom MATLAB code that will be made available upon acceptance of this manuscript.

## Authors Contributions

C.S. and A.Z. designed the research; C.S. and A.Z. performed the research; C.S., A.R. and A.Z. contributed analytic tools; C.S. and A.Z. analyzed the data; and C.S., A.R. and A.Z. wrote the paper.

## Acknowledgements

Fly, mouse and macaque icons in Fig. 6 and Supplementary Fig. 14 were obtained from the Mind the Graph platform (https://mindthegraph.com/). Human data were provided by the Human Connectome Project, WU-Minn Consortium (1U54MH091657; Principal Investigators David Van Essen and Kamil Ugurbil) funded by the 16 National Institutes of Health (NIH) institutes and centers that support the NIH Blueprint for Neuroscience Research, and by the McDonnell Center for Systems Neuroscience at Washington University. C.S. is funded by a Melbourne Research Scholarship. A.R. is funded by the Australian Research Council DECRA Fellowship (Ref: DE170100128). A.Z. is supported by the Australian National Health and Medical Research Council (NHMRC) Senior Research Fellowship B (1136649).

## SUPPLEMENTARY INFORMATION

### Note 1: Consistency of send-neutral-receiver classification of cortical regions across communication measures

Extending our analyses of correlations between send-receive asymmetries across communication measures, we compared the sender-neutral-receiver classifications of cortical regions obtained from navigation efficiency, diffusion efficiency and search information (Supplementary Table I). As expected by their mutual dependency on random walk processes, classifications for send-receive asymmetries under diffusion efficiency and search information were tightly related (76% accuracy, Supplementary Table Ic). While classification under navigation showed less agreement with the other measures (46% and 44% accuracy for diffusion and search information, respectively; Supplementary Table Ia,b), the obtained three-way classification accuracy remained larger than the 33% baseline expected by chance.

We further explored this relationship by considering two-way classifications of regions into sender or not sender, and receiver or not receiver. This allowed us to assess the statistical significance of the association between the classifications from two communication measures by using Fisher’s exact test. The significance of the obtained P-values further supports the classification consistency across communication measures (Supplementary Table Id-i).

### Note 2: Human connectome modularity and node participation

Modular decompositions were estimated for group-level connectomes computed as the average of all individual connectivity matrices for *N* = 256, 360, 512. The resulting group-level connectomes were thresholded at 10%, 15% and 20% connection density and analysed as weighted networks. Nodes were assigned to modules by means of the Louvain algorithm with iterative fine-tuning [1], as implemented in the Brain Connectivity Toolbox [2]. Briefly, the algorithm aims to identify partitions of the network that maximize within-module connection weights. This notion is formalized by the optimization of the modularity statistic

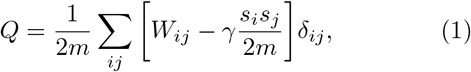

where *W_ij_* is the connection weight between nodes *i* and *j*, *γ* is the resolution parameter (set to default value of 1), *s_i_* is the strength of node *i*, 2*m* is a normalization constant equal to the sum of all connection weights in the network, and *δ_ij_* = 1 if *i* and *j* are assigned to the same module and 0 otherwise. For each group-level connectome, the Louvain algorithm was applied 100 times and the partition that maximized *Q* was selected. For each individual run of the algorithm, partitions were fine-tuned for *k* iterations until *Q_k_* – *Q*_*k*–1_ < 10^−5^ or *k* = 100. This procedure resulted in the identification of 7 modules for *N* = 256 (all connection densities); 6 modules for *N* = 360 (all connection densities); and 6, 7, 7 modules for *N* = 512, for 10%, 15%, 20% connection density, respectively.

The participant coefficient measures the diversity of inter-modular connections of individual nodes, and is defined as [3]

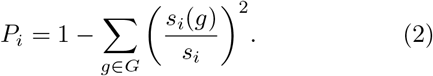

Here, *G* denotes the set of all modules, *s_i_* the strength of node *i* and *s_i_*(*g*) the strength of *i* within module *g*. High and low *P_i_* indicate that *i* has connections to nodes assigned to many and few distinct modules, respectively.

### Note 3: Randomized connectomes

We used three families of randomized connectomes: i) topologically randomized (rewired) networks, ii) topologically randomized cost-preserving networks and iii) spatially randomized (repositioned) networks. For *K* subjects, each individual connectome was used to generate ensembles of 1,000 surrogate networks for each family. Computing communication measures for these networks resulted in 1,000 sets of *C_N×N×K_* null communication efficiency matrices, per family, per communication measure. These sets were downsampled to subsystem resolution and used to compute the correlation between effective connectivity and null network send-receive communication asymmetries. Non-parametric P-values testing the hypothesis of a lack of difference between empirical and null correlations were computed as the proportion of times the communication asymmetry of null networks yielded stronger correlations than the one obtained for the empirical connectome.

Topologically randomized networks were computed using the Maslov-Sneppen rewiring routine [4] implemented in the Brain Connectivity Toolbox [2]. In this procedure, each connection was swapped between nodes once (on average), while maintaining the network’s original degree distribution and ensuring it remained connected.

One disadvantage of topologically randomized networks is the introduction of a disproportionate number of long-range connections, resulting in null networks with markedly increased wiring cost compared to the empirical network. To address this issue, we first computed the empirical wiring cost of a network as the sum of Euclidean distances between its connected nodes [5]. We then generated cost-preserving topologically randomized networks by adding a constraint to the Maslov-Sneppen routine, namely that connection swaps must not alter the original network’s cost by more than 1mm. Previous studies report that this leads to surrogate networks that match empirical wiring cost within a 0.1% error margin ^[6]^.

Spatially randomized networks are relevant for navigation, which routes information based on local knowledge of network geometry. They are constructed by randomly swapping the spatial positioning of nodes, while maintaining network topology unaltered.

### Note 4: Send-receive asymmetries and normality assumptions

Send-receive asymmetry between nodes/subsystems *i, j* is computed using a one-sample t-test on the distribution Δ(*i, j, k* = 1…*K*) = *C*(*i, j, k* = 1…*K*) – *C*(*j, i, k* = 1…*K*), where *C* denotes an asymmetric communication matrix and *K* is the number of subjects. A similar approach is used to compute regional asymmetries based on the node-level distributions *δ*(*i, k* = 1…*K*) (see *Materials and Methods, Send-receive communication asymmetry measures* for details). However, t-tests mandate an assumption of data normality. We tested the normality of δ for *i* = 1…360 and Δ for *i, j* = 1…*M*, where *M* = 7,17, 22 indicate the pairwise asymmetries at different cortical subsystems resolutions. Normality was assessed using a one-sample Kolmogorov-Smirnov test and we considered asymmetries computed for *N* = 360 at 15% connection density. At the regional level, the null hypothesis of normality for 87%, 47% and 85% of the δ distributions could not be rejected at the 5% significance level, for navigation, diffusion and search information, respectively. When accounting for the 360 multiple comparisons using Bonferroni correction, the respective percentages increased to 100%, 91% and 100%. For pairwise asymmetries between subsystems, the null hypothesis of normality could not be rejected at the 5% significance level for 100% of Δ distributions for *M* = 7,17, 22.

These results suggest that the majority of regional and pairwise communication asymmetries are normally distributed. However, to address possible deviations from normality, we recomputed send-receive asymmetries using a one-sample Wilcoxon rank-sum tests instead of the t-tests. Importantly, the Wilcoxon test is non-parametric and does not require variables to be normally distributed. The resulting w-statistic can be interpreted as an assessment of whether the median of a distribution is significantly smaller or greater than 0. We found that t-and w-statistic based send-receive asymmetries were highly correlated, at both regional and subsystem pairwise levels and for all connection density thresholds. Regional send-receive asymmetries (for *N* = 360) under navigation: Pearson correlation coefficient *r* = 0.96,0.97,0.97 for 10%, 15% and 20% connection density thresholds, respectively; under diffusion: *r* = 0.68 for all connection densities; and under search information: *r* = 0.89 for all connection densities. Pairwise subsystem send-receive asymmetries (for *N* = 360 and 15% connection density) under navigation: *r* = 0.96,0.95 for *M* = 17, 22, respectively; under diffusion: *r* = 0.87,0.89 for *M* = 17, 22, respectively; and under search information: *r* = 0.87,0.88 for *M* = 17, 22, respectively. Accordingly, we found that t- and w-statistic send-receive communication asymmetries were consistently correlated with the directionality of effective connectivity (Supplementary Fig. 10a,b).

### Note 5: Send-receive asymmetries for connectomes mapped with probabilistic tractography

We sought to determine whether the send-receive asymmetry of the human connectome is robust to alternative connectome reconstruction techniques. To this end, we recomputed the connectomes of the same *K* = 200 subjects using a probabilistic tractography pipeline. Using MRtrix3 [7], we applied multi-shell, multi-tissue constrained spherical deconvolution to estimate the fibre orientation distribution of each white matter voxel [8]. Probabilistic tractography was carried out with the following parameters: iFOD2 algorithm [9], 5 × 10^6^ streamlines, 0.5 mm tracking step-size, 400 mm maximum streamline length and 0.1 fractional anisotropy cutoff for streamline termination. The resulting tractograms were used to construct human connectomes comprising *N* = 256, 360, 512 cortical regions. The obtained connectivity matrices were thresholded at 10%, 15% and 20% connection density and analysed as weighted networks. Regional and subsystem level send-receive communication asymmetries were computed as for the deterministic connectomes (*Materials and Methods, Send-receive communication asymmetry measures*).

We found that deterministic and probabilistic send-receive asymmetries were strongly correlated for all communication measures, connection density thresholds, parcellation resolutions and cortical subsystem partitions (Supplementary Table II and Supplementary Fig. 11). For instance, deterministic and probabilistic regional asymmetries showed a Pearson correlation coefficient of *r* = 0.75,0.74, 0.77 under navigation, diffusion and search information, respectively (*N* = 360 at 15% connection density). Similarly, pairwise asymmetries between the two tractography approaches for *M* = 22 subsystems showed *r* = 0.70,0.76,0.76 under navigation, diffusion and search information, respectively (*N* = 360 at 15% connection density). All *P* < 10^−30^.

Communication asymmetries derived from probabilistic connectomes were also correlated with effective connectivity directionality (Supplementary Fig. 10c,d; e.g., *r* = 0.64, 0.48, 0.45 for *M* =17 under navigation, diffusion and search information, respectively). Interestingly, in most cases, probabilistic asymmetries yielded stronger correlations to effective connectivity directionality than the original deterministic ones. In addition, probabilistic asymmetries did not lead to the drop in association strength for *M* = 17 in fMRI session 2 present in deterministic-based results. However, regional probabilistic-based asymmetries were not associated to the cortical gradient of functional heterogeneity (*P* > 0.05 across all communication measures and parcellation resolutions).

Taken together, these results indicate that send-receive communication asymmetry measures are robust to different tractography pipelines. Regional and pairwise send-receive asymmetries were strongly correlated across deterministic and probabilistic connectome reconstructions and both approaches yielded strong correlations with neural signalling directionality estimated with spectral DCM. Differences in the association strength with effective connectivity directionality and local functional heterogeneity suggest the possibility that deterministic and probabilistic connectomes are better able to capture, respectively, regional and pairwise resting-state functional activity.

### Note 6: Senders, receivers and the subcortex

In the main manuscript, we analysed brain networks comprising exclusively of cortico-cortical connections due to challenges in mapping connectomes containing subcortical structures. However, subcortical regions are important mediators of signalling between cortical areas. To investigate the potential impact of subcortical structures to our results, we aimed to reproduce our send-receive asymmetry findings in connectomes including the subcortex. To this end, we mapped the connectomes of the same *K* = 200 subjects using a gray matter parcellation comprising 360 cortical and 14 subcortical regions (left and right thalamus, caudate, putamen, pallidum, hippocampus, amygdala and accumbens-area). The 360 cortical areas were defined according to the HCP MMP1.0 atlas [10], while subcortical structures were derived from the Freesurfer parcellation provided by the HCP (*aparc+aseg* file).

An important challenge in mapping whole-brain connectomes including the subcortex is that certain subcortical structures, in particular the thalamus, are composed of both gray and white matter, functioning both as signal relays and information processing units [11]. However, current parcellations of the subcortex do not provide sufficient detail to differentiate between gray matter nuclei and white matter fibers within subcortical structures. In order to address this issue, we employed a data-driven tractography pipeline with respect to subcortical structures. Subcortical regions were included in both white and gray matter masks. During tractography, subcortical voxels with sub-threshold fractional anisotropy (FA) were interpreted as gray matter, while supra-threshold voxels were considered white matter. Hence, if a streamline tracking through a subcortical region encounters a voxel with low FA (< 0. 1), this is interpreted as a portion of gray matter within the subcortical structure, and the streamline’s endpoint is assigned to the current subcortical region. However, if a streamline enters and exits a subcortical structure without passing through low FA voxels, this is interpreted as a white matter tract connecting two other (cortical or subcortical) regions. In this case, the streamline endpoint is not assigned to the current subcortical region, providing a data-driven model of subcortical structures as signal relays.

Using this approach, we applied the deterministic tractography algorithm described in *Materials and Methods, Human Connectomes* to map connectivity matrices comprising *N* = 360+14 = 374 nodes. Previously, in order to account for biases towards higher connectivity strength in larger regions, streamline counts between pair of regions were normalized by the product of their surface areas. In this case, since subcortical regions were defined only in volume space, we instead used the pairwise product of regional volumes as a normalization factor. Cortical surface-based parcellations were registered to subject-specific T1-weighted images and the number of voxels comprising gray matter regions was used as an approximation of their volume. This resulted in *K* = 200 subject-level 374 × 374 weighted connectivity matrices, which were thresholded at 10%, 15% and 20% connection density and analysed as weighted networks. It is noteworthy that subcortical regions ranked amongst the highest-degree nodes in the obtained connectomes (for the average degree across subjects, the thalamus, putamen, hippocampus, pallidum, amygdala, accumbens-area and caudate, ranked, respectively, amongst the top 0.83%, 1.9%, 3.6%, 4.4%, 6.7%, 8.6% and 11.4% most connected nodes).

We began by investigating whether cortical send-receive asymmetries computed on cortico-cortical connectomes were comparable to those derived from connectomes including the subcortex. At the regional level, we found that send-receive asymmetries of cortical regions were consistent across the two sets of connectomes (Supplementary Fig. 12). Pearson correlation coefficients between cortical send-receive asymmetries for connectomes with and without the subcortex: *r* = 0.86,0.84,0.85 under navigation; *r* = 0.92,0.93,0.93 under diffusion; and *r* = 0.90,0.90,0.90 under search information, for 10%, 15% and 20% connection density, respectively (all *P* < 10^−96^; see Supplementary Fig. 12a,e,i for scatter plots of these associations at 15% connection density). Accordingly, the classification of cortical regions into senders, neutral and receivers was robust to the inclusion of the subcortex: three-way classification accuracy of 69%, 74% and 72% for asymmetries under navigation, diffusion and search information, respectively, at 15% connection density. After the inclusion of subcortical regions, primary sensory regions remained senders (with the exception of V1 for navigation and the right hemisphere M1 for search information), while portions of the precuneus, frontal and prefrontal cortices remained as receivers (Supplementary Fig. 12b,f,j).

Despite these consistencies, the inclusion of subcortical structures did influence the send-receive asymmetry of cortical regions. Supplementary Fig. 12c,g,k shows the difference in regional asymmetries for connectomes with and without the subcortex. Positive differences (shown in red) indicate shifts towards outgoing communication efficiency (increase in send-receive asymmetry), while negative differences (shown in blue) indicate shifts towards incoming communication efficiency (decrease in send-receive asymmetry). In other words, following the inclusion of the subcortex, regions shown in red and blue became relatively more biased towards sending and receiving information, respectively. For navigation (Supplementary Fig. 12c), considering subcortical communication pathways reduced the propensity of visual, somatosensory and auditory cortices towards outgoing communication efficiency. Accordingly, portions of the temporal-parietal-occipital junction, MT+ complex, and medial and dorsolateral cortices had their propensity towards incoming communication efficiency attenuated. Similar patterns of asymmetry differences were observed under diffusion and search information (Supplementary Fig. 12g,k), with the marked distinction that early visual areas (V1 and V2) increased their propensity towards outgoing communication. Importantly, as described above, the majority of cortical regions were consistently classified as senders, neutral or receivers, regardless of whether subcortical structures were present. Hence, these results indicate that subcortical nodes, by mediating communication between cortical areas, influence the intensity (rather than the sign) of cortical send-receive communication asymmetries.

Next, we turned our attention to the send-receive asymmetry of individual subcortical regions. Across all communication measures, all subcortical structures showed a marked bias towards outgoing communication efficiency (Supplementary Fig. 12d,h,l). This is an interesting result, particularly because it opposes the propensity of high-degree regions towards incoming communication reported for diffusion and search information (see *Results, Senders and receivers of the human connectome*). The consistent classification of subcortical regions as senders (with exception of the accumbens-area) potentially reflects their role as inputs of sensory signals to primary cortices.

We also sought to investigate pairwise asymmetries between the subcortex and cortical subsystems. To this end, subcortical regions were grouped into a single subsystem. In keeping with regional results, pairwise asymmetry matrices between cortical subsystems were robust to the inclusion of the subcortex (Supplementary Fig. 13; navigation: *r* = 0.89, 0.88, diffusion: *r* = 0.69, 0.76, search information: *r* = 0.71, 0.80, for *M* = 7, 22, respectively. All *P* < 10^−3^). Accordingly, individual cortical subsystems showed similar classification as senders and receivers following the addition of subcortical regions (Supplementary Fig. 13b,e,h,l,o,r). Interestingly, a comparison of the send-receive asymmetry of cortical subsystems with (colored horizontal bars) and without the subcortex (colored circles) corroborates the notion that subcortical pathways generally attenuate the communication asymmetry between cortical systems, but do not interfere with their directionality. Once more in agreement with regional results, the subcortex subsystem was consistently classified as a prominent sender.

Finally, we found that send-receive asymmetries between cortical subsystems remained correlated to effective connectivity following the inclusion of the subcortex (Supplementary Fig. 10e,f). Interestingly, associations for *M* = 17 subsystems (Yeo resting-state functional modules [12]) were weakened, while associations for *M* = 22 (HCP contiguous modules [10]) were strengthened, with search information asymmetry leading to correlations as high as *r* = 0.68 (resting-state session 2, 15% connection density).

### Note 7: Senders and receivers of non-human connectomes

In this section, we explore send-receive asymmetries computed on symmetrized (undirected) non-human connectomes *(Materials and methods, Symmetrized nonhuman connectomes).* For sake of conciseness, we refer to those as simply send-receive asymmetries.

We begin by noting that for all non-human connectomes considered, send-receive asymmetries under navigation were not correlated to send-receive asymmetries under diffusion or search information (all *P* > 0.05). As with the human results, diffusion and search information asymmetries were strongly correlated across species (*r* = 0.96,0.75,0.75 for fly, mouse and macaque regional send-receive asymmetries, respectively). Hence, the obtained classification of regions as senders and receivers was different between navigation and random walk based measures. A possible explanation for this disparity lies in the high connection density of the non-human connectomes (fly: 83%, 89%, mouse: 53%, 70%, macaque: 66%, 79%, directed (original) and undirected (symmetrized) connection densities, respectively). In undirected networks (as obtained after connectome symmetrization), there is no navigation asymmetry between directly connected node pairs (if *W*(*i,j*) = *W*(*j,i*) ≠ 0 it follows that *E_nav_*(*i,j*) = *E_nav_*(*j, i*). See *Materials and Methods, Navigation efficiency*). This may lead to a decrease in the biological relevance of navigation asymmetries in connectomes for which most regions are directly connected. Meanwhile, the stochastic nature of communication measures based on random walks gives rise to send-receive asymmetries even in densely connected networks. In support of this notion, we found that senders and receivers identified under diffusion and search information were consistent with the putative functional roles of different brain regions in the fly, mouse and macaque. Moreover, send-receive asymmetries computed on binary connectomes yield the most biologically relevant classification of senders and receivers. Given the agreement between diffusion and search information, we focus on describing the results obtained for binary diffusion send-receive asymmetry in the following paragraphs.

A limitation of non-human connectome analyses is the lack of subject-level data. Importantly, this shortcoming precludes the use of the same send-receive asymmetry framework developed for humans. As a result, send-receive asymmetry for these species are not defined statistically at the level of individual regions or pairs of regions. To address this limitation, we partitioned the nodes comprising non-human connectomes into *M* previously defined subsystems of the fly, mouse and macaque nervous systems. We downsampled the node-level *N × N* send-receive asymmetry matrices into subsystem-level *M* × *M* matrices. As a result, each subsystem pair is associated to a distribution of node-level pairwise asymmetries. For each subsystem pair *i, j*, we computed whether the mean of their distribution of node-level pairwise asymmetries was significantly larger than 0 by means of a one-sample t-test. Send-receive asymmetry between subsystems was defined as the resulting t-statistic. Note that while human asymmetries were statistically defined based on cross-subject distributions, here we define non-human subsystem asymmetries across node pairs. This allowed us to statistically test hypotheses on the agreement between communication asymmetry and the putative functional roles of subsystems of non-human connectomes.

Supplementary Fig. 14a shows the regional send-receive diffusion asymmetry of the 49 neuronal populations comprising the fly connectome. Following previ ous work on decentralized communication in the fly connectome, we classified neuronal populations into three groups: (i) sensors (ii) effectors and (iii) others [13]. Sensor nodes are involved in the processing and transducing of sensory information, and are thus hypothesised to be senders. Effectors are conjectured to integrate signals from different neuronal populations in order to coordinate motor execution, and are thus hypothesised to be receivers. In line with these hypotheses, 4 out of 6 sensor nodes were classified as senders (positive send-receive asymmetry) while all effector nodes were classified as receivers (negative send-receive asymmetry). Additionally, we found that the mean sensor ! effector and other ! effector asymmetries were significantly larger than 0 (*P* =1 × 10^−9^, 5 × 10^−24^, respectively. Supplementary Fig. 14b). The average sensor ! other asymmetry was positive, but not significantly larger than 0 (*P* = 0.11). Asymmetries between sensor ! effector and other ! effector nodes were not statistically different (two-sample t-test *P* = 0.46). Together, these findings indicate that diffusion send-receive asymmetry recapitulates the putative functional roles of neural populations in the fly connectome by characterizing sensor nodes as senders and effector nodes as receivers (Supplementary Fig. 14c).

Supplementary Fig. 14d shows the regional send-receive diffusion asymmetry of regions comprising the mouse connectome (values were averaged across homotopic regions). Consistent with the human results, we found that primary auditory (AUDp) and visual (VISp) areas were senders (positive send-receive asymmetry). However, primary motor (MOs) and sensory (SSp) were classified as receivers. We partitioned brain regions into 8 modules according to a previously established modular decomposition of the mouse connectome [14]. The 8 modules are: somatosensory-motor (SS-M), brainstem-cerebellum (BS-CE), auditory (AU), visual (VI), olfactory (OL), hippocampal (HI), hypothalamic (HY), and high-participation (Hi-Par, i.e., regions characterized by a large proportion of intermodule connections that could not be consistently assigned to a single module). Generally, subsystem associated with the processing of sensory information were senders (olfactory, visual and auditory), with the exception of somatosensory-motor (Supplementary Fig. 14e,f). The brainstem-cerebellum subsystem was the most prominent sender, potentially reflecting the role of subcortical structures in relaying sensory signals to primary cortices. In agreement with the human findings, regions with diverse intermodule connectivity (high participation) were prominent receivers. Together, these results indicate that the undirected topology of the mouse connectome leads to biases in outgoing information for auditory, visual and olfactory regions. Meanwhile, sensory and motor areas may be more dependent on axonal directionality to propagate information to higher-order areas of the mouse cortex.

Finally, Supplementary Fig. 14g shows the regional send-receive diffusion asymmetry of the 29 regions comprised in the macaque connectome. In agreement with results reported for the human connectome, areas of the sensory (area 2), visual (V1 and V2) and motor (F1 and ProM) cortices were classified as senders (positive send-receive asymmetry), while portions of the frontal (8m and 8l) and prefrontal cortices (46d, 9/46v, 9/46d) were classified as receivers (negative send-receive asymmetry). Supplementary Fig. 15 compares the cortical projections of human and macaque send-receive asymmetries and provides further evidence for consistencies between the results of the two species: expanses of the occipital cortex and sensory-motor strip are classified as senders, while portions of the prefrontal cortex are classified as receivers. Certain differences between the two species were observed: higher-order visual areas such as V4 were receivers in humans but senders in the macaque, while expanses of the temporal lobe appear as senders in humans and receivers in the macaque. Cortical regions were assigned to 6 modules according to a previously established partition of the macaque connectome [15]. Occipital and prefrontal modules were the most prominent senders and receivers, respectively (Supplementary Fig. 14h,i). Taken together, these findings indicate that the interaction between decentralized network communication measures and connectome topology leads to send-receive asymmetries that recapitulate uni-to heteromodal cortical hierarchies in both human and macaque connectomes. These results suggest that topological properties of undirected connectomes contributing to neural signalling directionality are phylogenetically conserved across higher primates.

It is important to notice that these analyses constitute a first account of send-receive asymmetry in nonhuman connectomes. Further work is necessary to consolidate the results presented here. Potential future steps in this direction include (i) a conceptualization of statistically defined send-receive asymmetry inferred from single-subject networks and (ii) a better understanding motivating the observed biological relevance of binary diffusion-based communication asymmetries.

---

**TABLE I.**
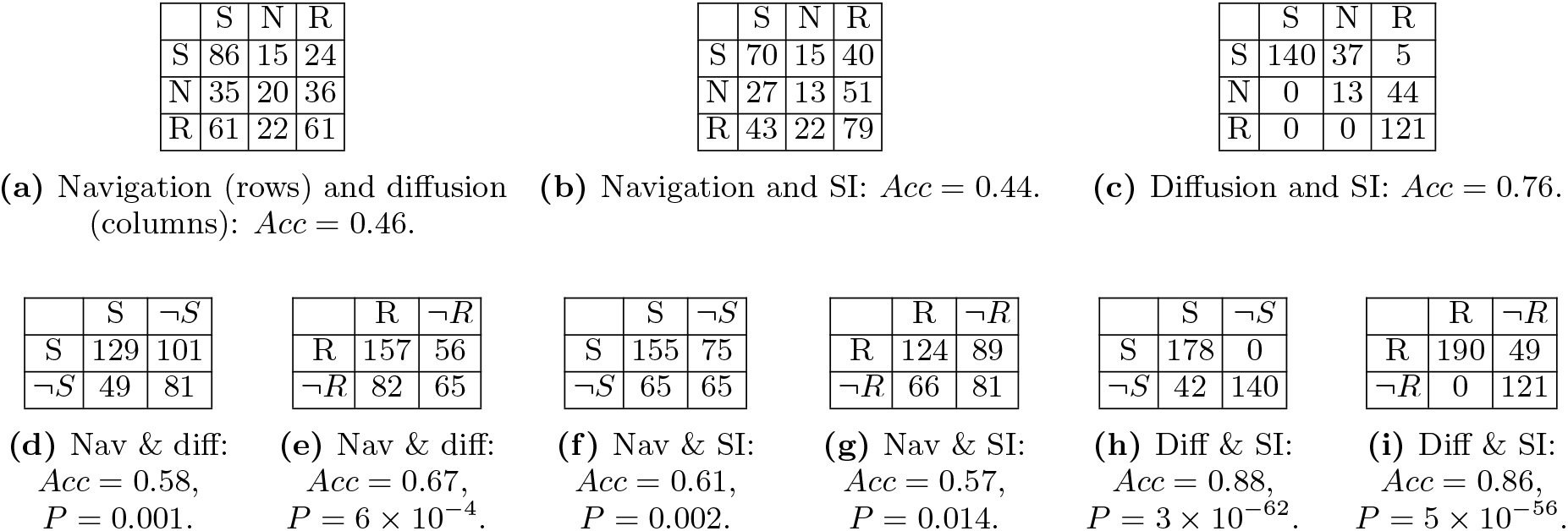
Comparison of the classification of cortical regions as senders (S), neutral (N) and receivers (R) across network communication measures. The classification accuracy (*Acc*) is computed as the sum of values in the main diagonal (number of consistently classified regions) divided by the sum of values in the table (total number of regions). **(a-c)** Three-way contingency tables of the classification obtained from the send-receive asymmetries of two communication measures. Measures listed first and second in the captions have their classes displayed in the rows and columns of the tables, respectively. **(d-i)** Two-way contingency tables of the classification obtained from the send-receive asymmetries of two communication measures. In this case, for each pair of measures, regions are classified as sender (S) or not sender (*S*¬), and receiver (R) or not receiver (¬*R*). P-values obtained from Fisher’s exact test were used to examine the significance of the association between the classifications of two measures.

**TABLE II.** Pearson correlation coefficients between deterministic and probabilistic send-receive asymmetries for different communication measures, connection density (CD) thresholds and parcellation resolutions. All associated *P* <10^−30^.

| Parcellation resolution | Navigation |  |  | Diffusion |  |  | Search Info |  |  |
| --- | --- | --- | --- | --- | --- | --- | --- | --- | --- |
|  | 10% CD | 15% CD | 20% CD | 10% | 15% | 20% | 10% | 15% | 20% |
| 256 | 0.75 | 0.71 | 0.68 | 0.71 | 0.72 | 0.71 | 0.74 | 0.74 | 0.75 |
| 360 | 0.77 | 0.75 | 0.73 | 0.75 | 0.74 | 0.74 | 0.77 | 0.77 | 0.77 |
| 512 | 0.74 | 0.72 | 0.70 | 0.67 | 0.66 | 0.65 | 0.65 | 0.76 | 0.76 |

**Supplementary Fig. 1.**
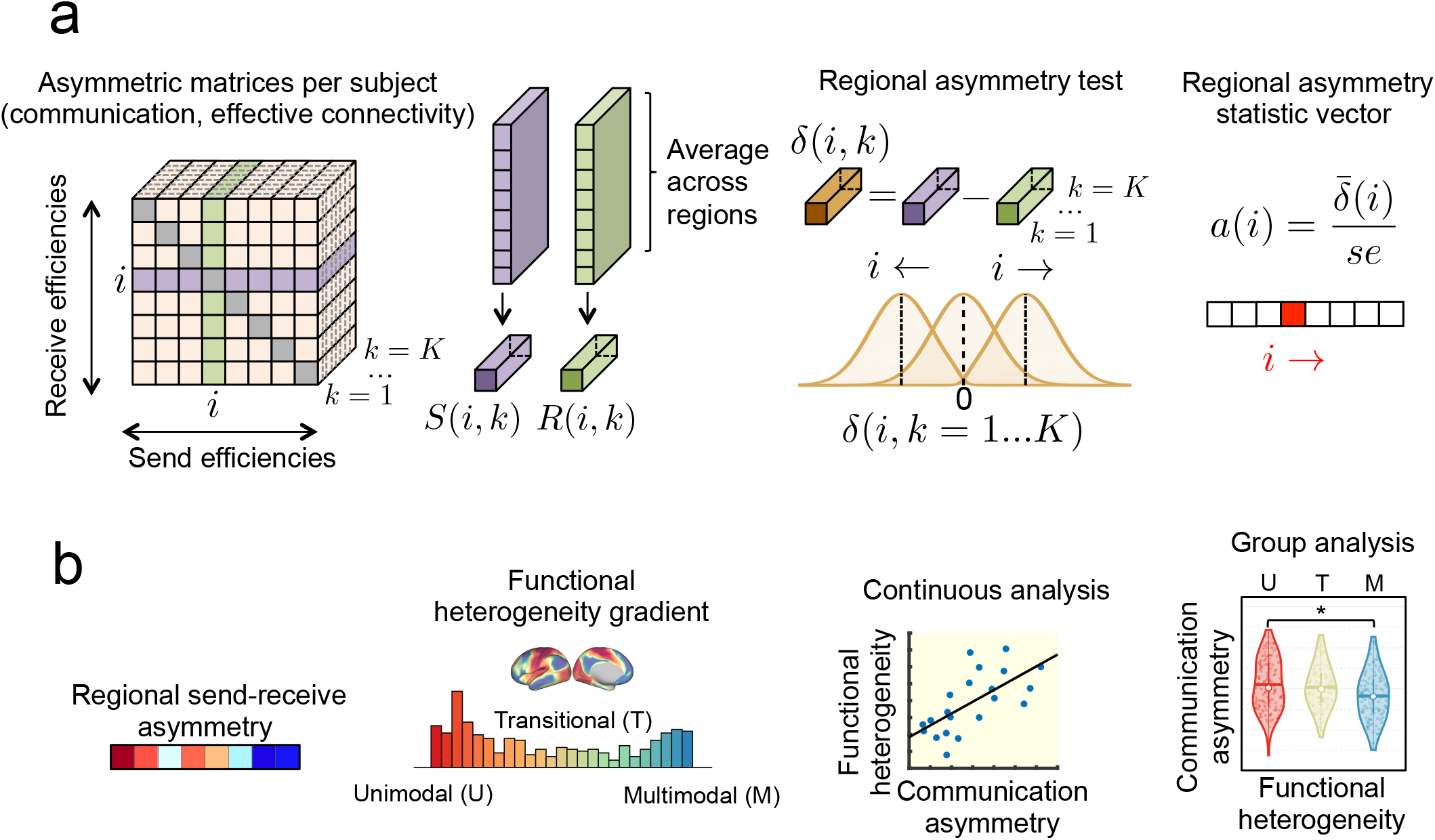
Methodology overview of regional send-receive asymmetry analyses. **(a)** Schematic of the regional communication asymmetry test as described in *Materials and Methods, Send-receive communication asymmetry measures*. **(b)** Schematic of the comparison between regional send-receive asymmetry and the Margulies’ gradient of functional heterogeneity [16]. Cortical regions are divided into unimodal, transitional and multimodal based on their placement along the functional gradient, as per described in *Materials and Methods, Cortical gradient of functional heterogeneity*. Comparisons between send-receive asymmetry and functional heterogeneity are performed by means of continuous (i.e., linear correlation) and group-wise (i.e., between group differences) analyses.

**Supplementary Fig. 2.**
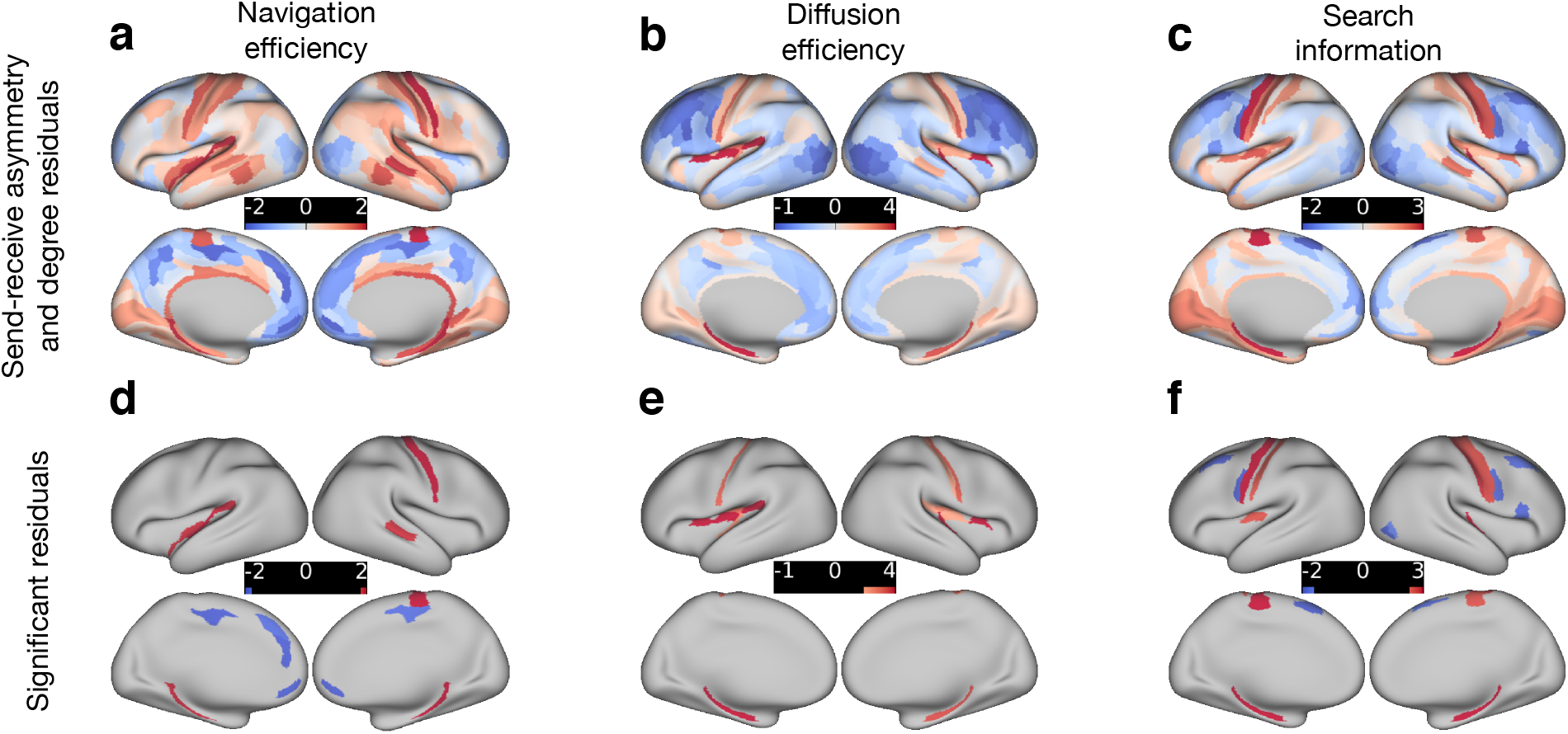
Standardized residuals 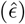 obtained from regressing out node degree from send-receive asymmetry (*N* = 360 at 15% connection density) under **(a,d)** navigation, **(b,e)** diffusion, and **(c,f)** search information. Regions shown in red 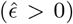 and blue 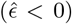 are, respectively, stronger senders and receivers than expected based on their degree alone. Regions with statistically significant residuals 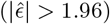 are highlighted in the bottom row.

**Supplementary Fig. 3.**
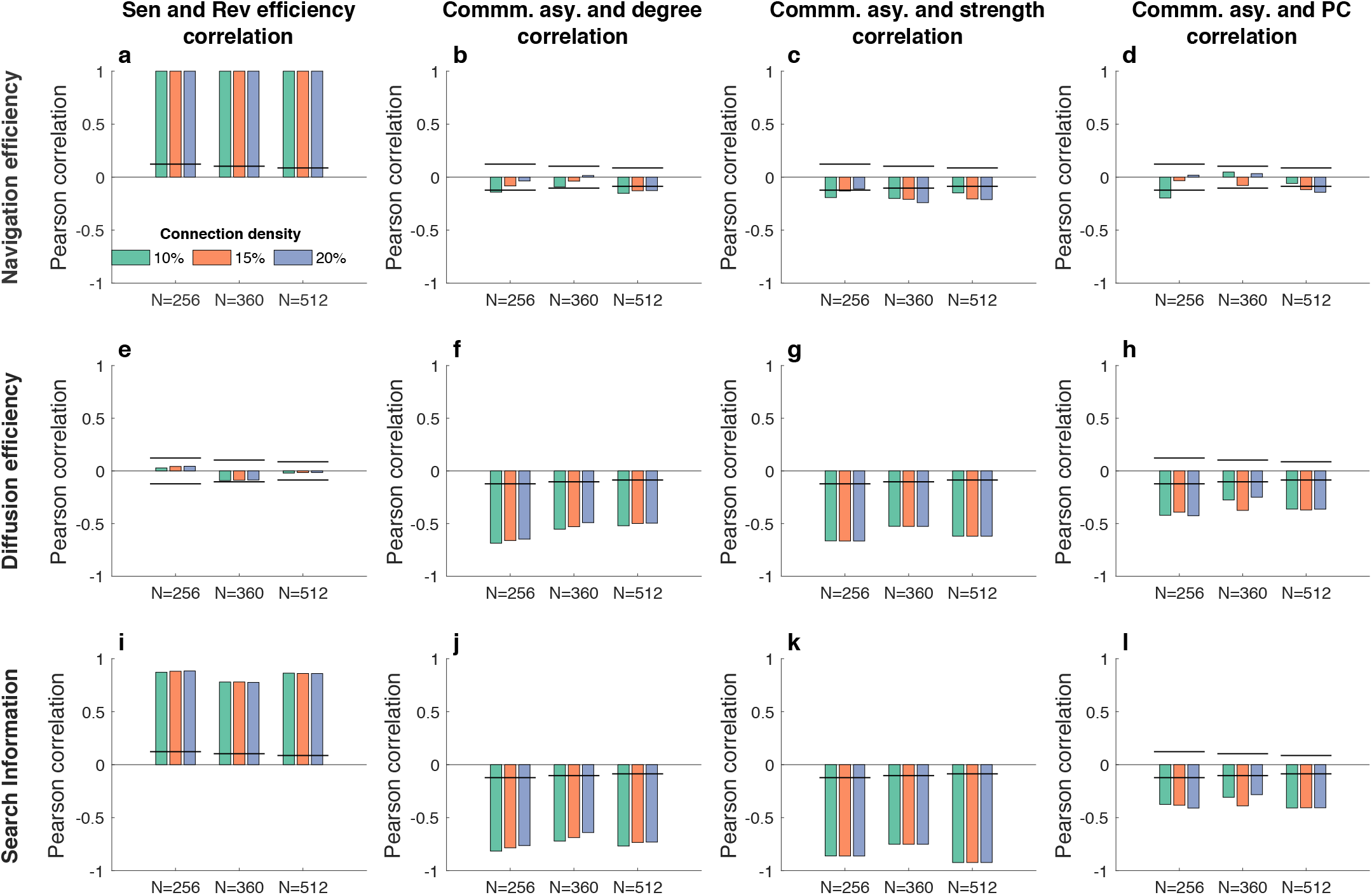
Replication analyses regarding sending efficiency, receiving efficiency and send-receive communication asymmetry for *N* =256, 360, 512 parcellation resolutions and 10%, 15% and 20% connection density thresholds. Vertical axes indicate the Pearson correlation’s *r*, while black horizontal lines mark the effect size correspondent to a correlation with *P* =0.05 for each *N*. **(a)** Correlation between sending and receiving navigation efficiencies. **(b)** Correlation between regional navigation asymmetry and node degree (averaged across the connectomes of all participants). **(c)** Same as b, but for node strength. **(d)** Same as a, but for node participation coefficient. **(e-h)** Same as a-d, but for diffusion efficiency. **(i-l)** Same as a-d, but for search information.

**Supplementary Fig. 4.**
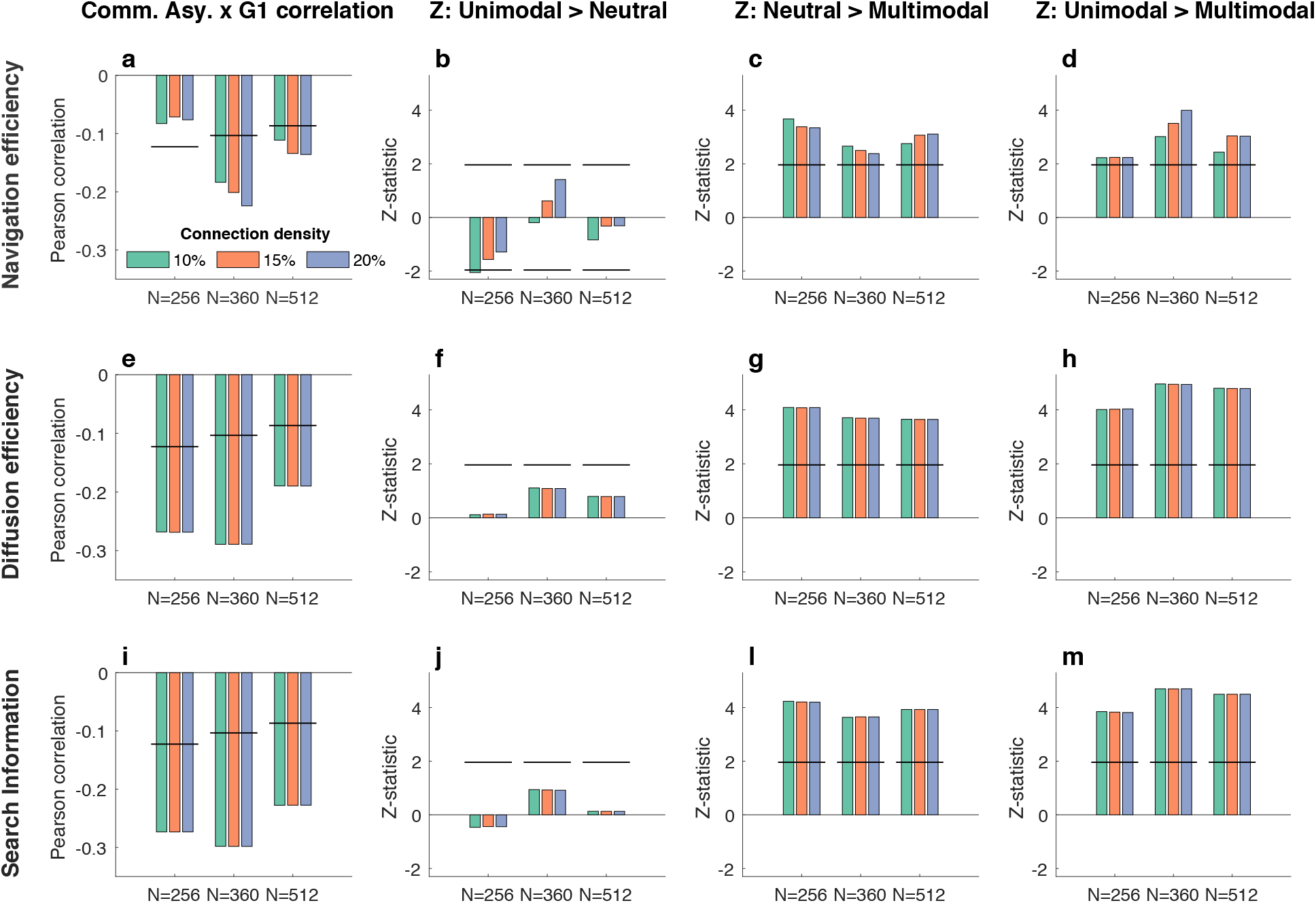
Replication analyses regarding the relationship between communication asymmetry and functional heterogeneity for *N* =256, 360, 512 parcellation resolutions and 10%, 15% and 20% connection density thresholds. The obtained results are consistent across parcellations and connection densities, with the exception of the lack of correlation between send-receive asymmetry and the gradient of functional heterogeneity for *N* =256. **(a)** Pearson correlation between regional navigation asymmetry and functional heterogeneity. Black horizontal lines mark the effect size correspondent to a correlation with *P* =0.05 for each *N*. **(b)** Z-statistic from a two-sided Wilcoxon test evaluating the hypothesis that the median navigation asymmetry of unimodal regions is larger than that of neutral regions. Black horizontal lines mark the value of a Z-statistic correspondent to *P* = 0.05. **(c)** Same as b, but for neutral and multimodal regions. **(d)** Same as b, but for unimodal and multimodal regions. **(e-h)** Same as a-d, but for diffusion efficiency. **(i-m)** Same as a-d, but for search information.

**Supplementary Fig. 5.**
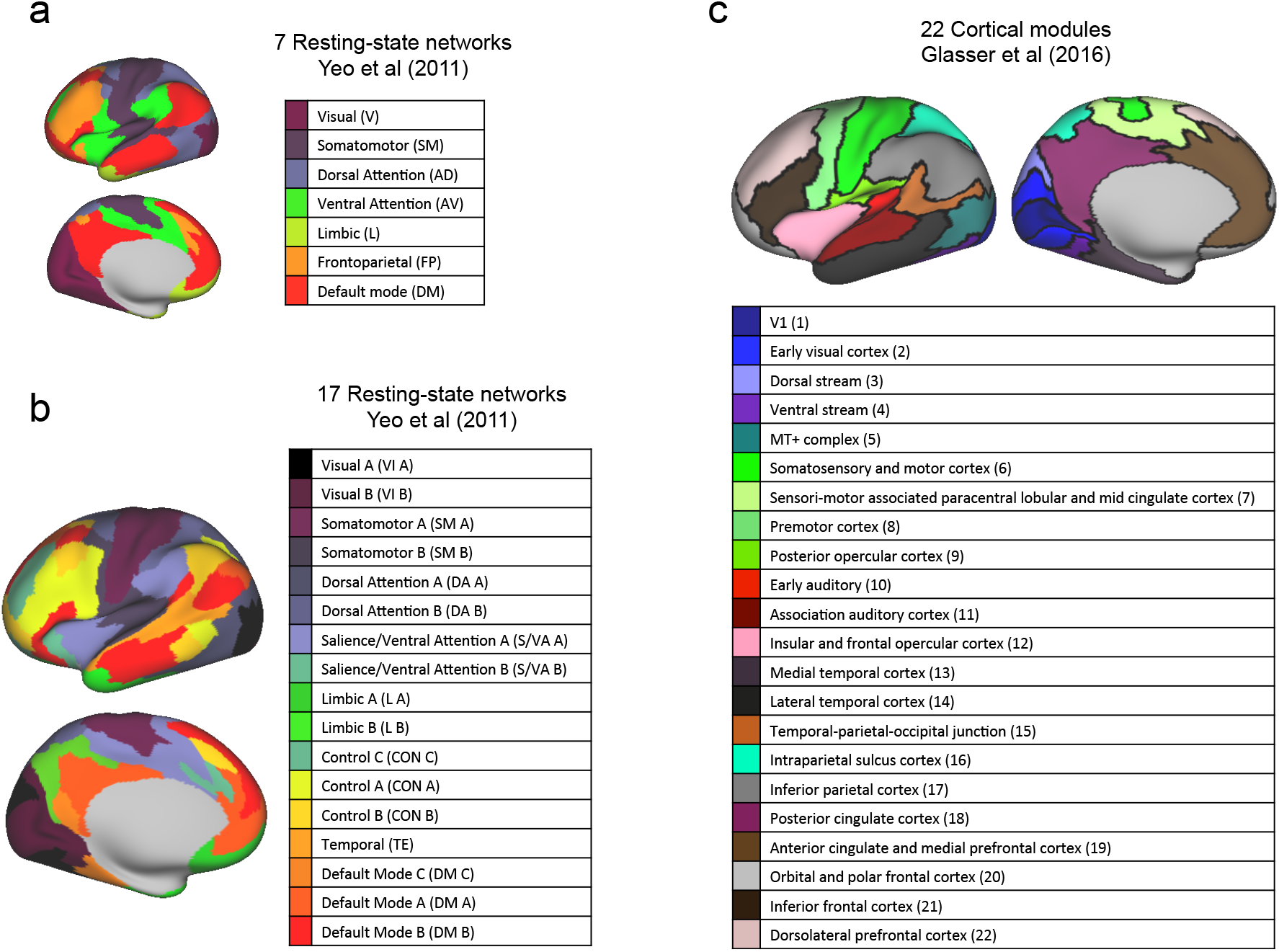
Definition of the *M* = 7, 17, 22 cortical subsystems utilized in sections *Send-receive communication asymmetries of cortical subsystems Send-receive communication asymmetry and effective connectivity*.

**Supplementary Fig. 6.**
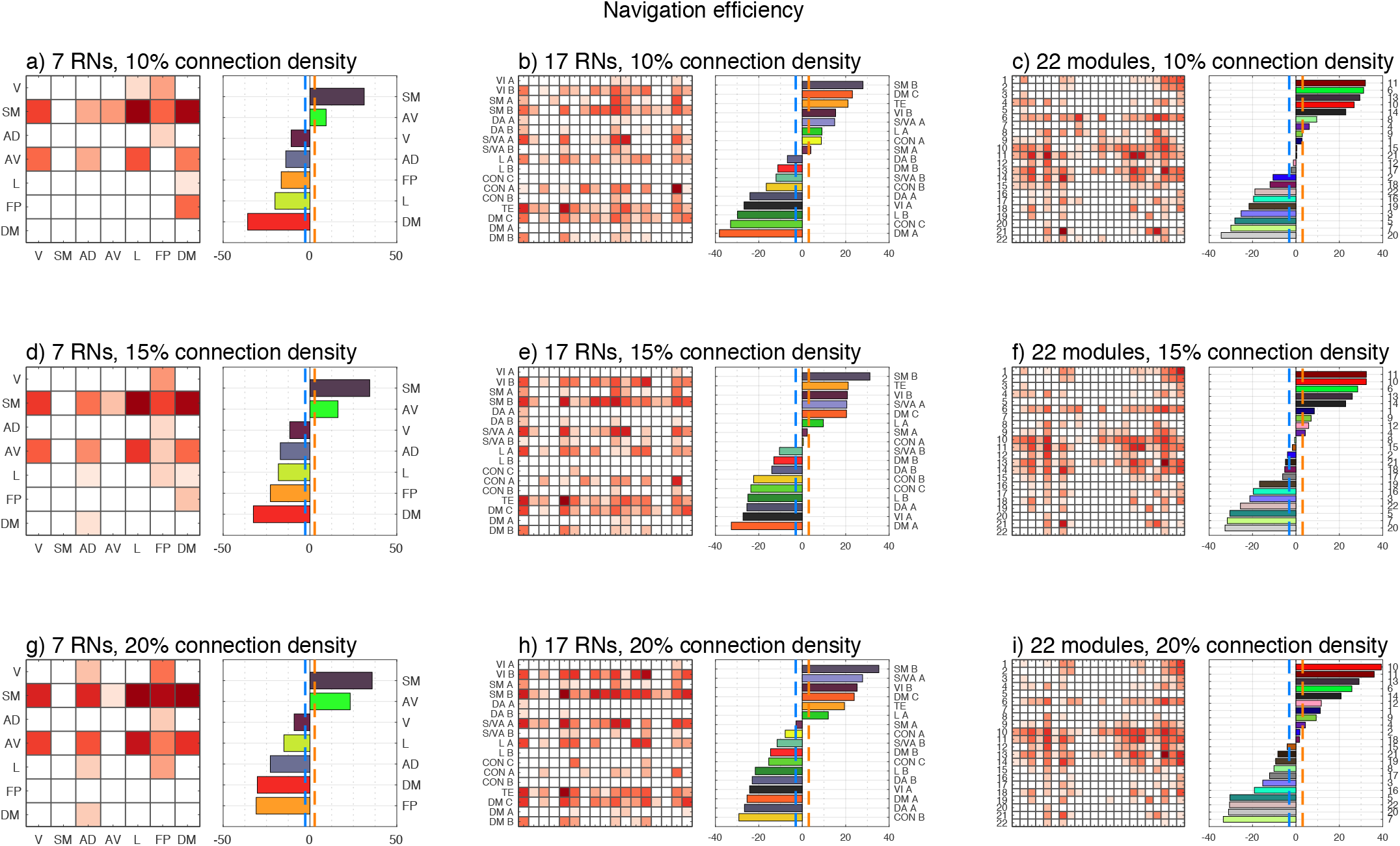
Send-receive navigation asymmetry of cortical subsystems for *M* = 7, 17, 22 and 10%, 15% and 20% connection density thresholds. Send-receive asymmetry matrices were thresholded to display only statistically significant values, while accounting for multiple comparisons. For ease of visualization and without loss of information (since *A*(*i, j*) = −*A*(*j, i*)), negative values were omitted. Thus, *A*(*i, j*) > 0 denotes that communication takes place more efficiently from *i* to *j* than from *j* to *i*.

**Supplementary Fig. 7.**
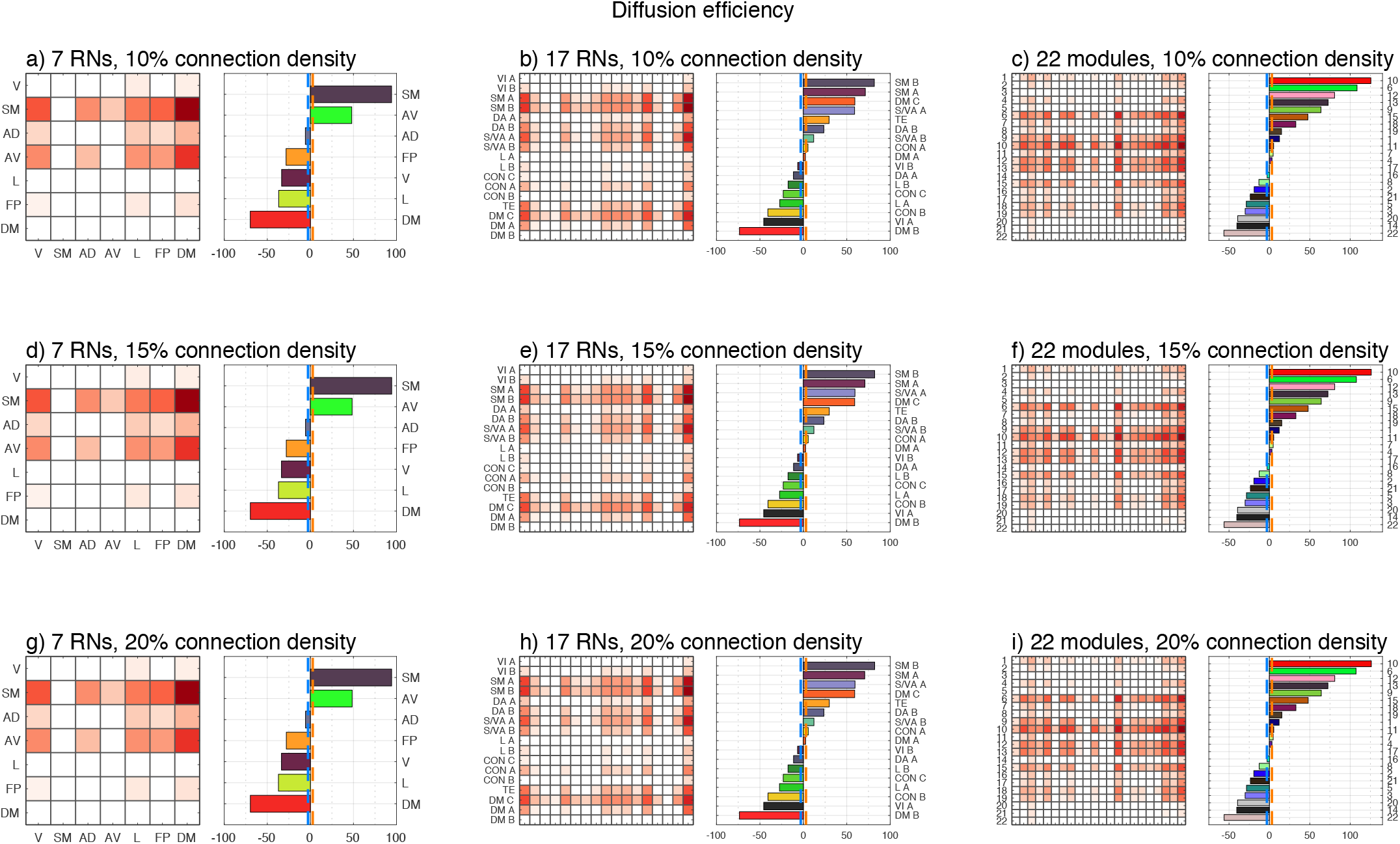
Send-receive diffusion asymmetry of cortical subsystems for *M* = 7, 17, 22 and 10%, 15% and 20% connection density thresholds. Send-receive asymmetry matrices were thresholded to display only statistically significant values, while accounting for multiple comparisons. For ease of visualization and without loss of information (since *A*(*i, j*) = −*A*(*j, i*)), negative values were omitted. Thus, *A*(*i, j*) > 0 denotes that communication takes place more efficiently from *i* to *j* than from *j* to *i*.

**Supplementary Fig. 8.**
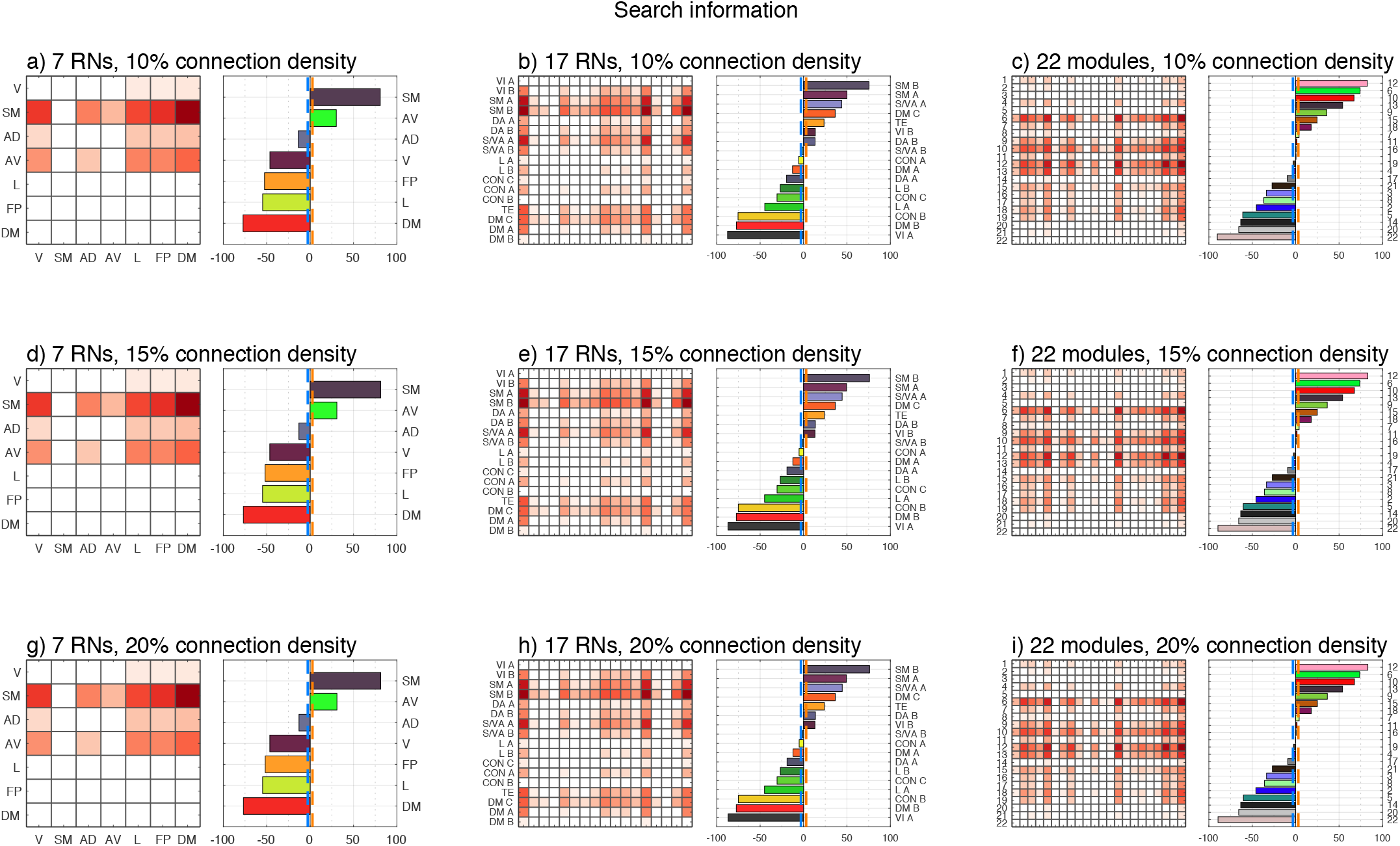
Send-receive search information asymmetry of cortical subsystems for *M* = 7, 17, 22 and 10%, 15% and 20% connection density thresholds. Send-receive asymmetry matrices were thresholded to display only statistically significant values, while accounting for multiple comparisons. For ease of visualization and without loss of information (since *A*(*i, j*) = −*A*(*j, i*)), negative values were omitted. Thus, *A*(*i, j*) > 0 denotes that communication takes place more efficiently from *i* to *j* than from *j* to *i*.

**Supplementary Fig. 9.**
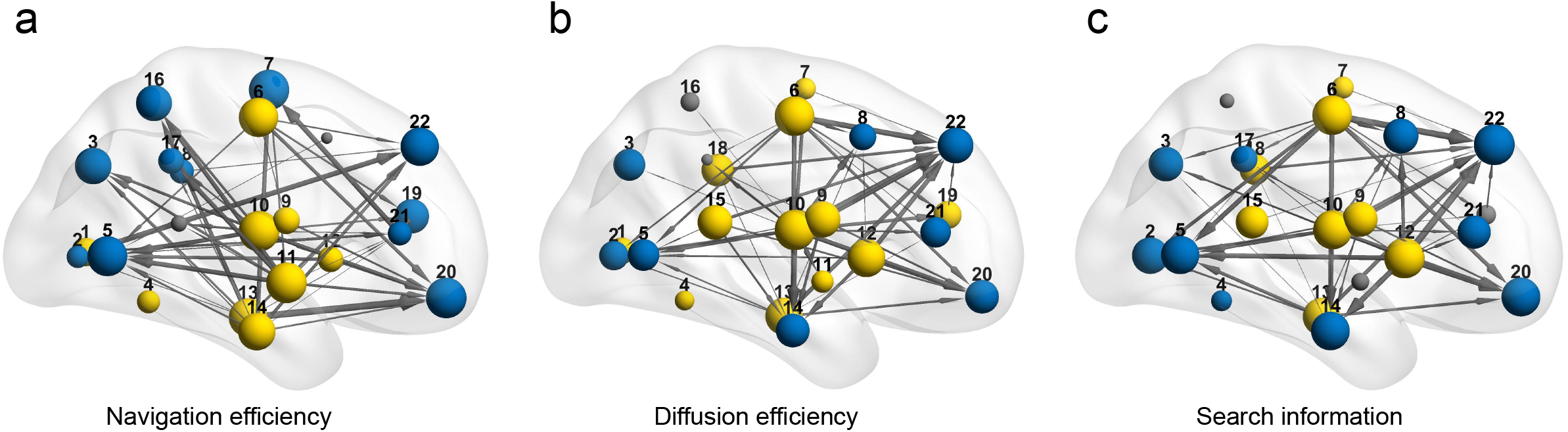
Network visualization of the 10% strongest pairwise send-receive asymmetries between *M* = 22 cortical subsystems (*N* = 360 at 15% structural connection density threshold) under **(a)** navigation, **(b)** diffusion and **(c)** search information. A directed connection between from subsystem *i* to subsystem *j* indicates that the communication efficiency from *i* to *j* is significantly higher than the communication efficiency from *j* to *i*. Connection width is proportional to the strength of the send-receive asymmetry. Subsystems classified as senders, neutral and receivers are shown in yellow, gray and blue, respectively. Visualization developed with BrainNet Viewer [17].

**Supplementary Fig. 10.**
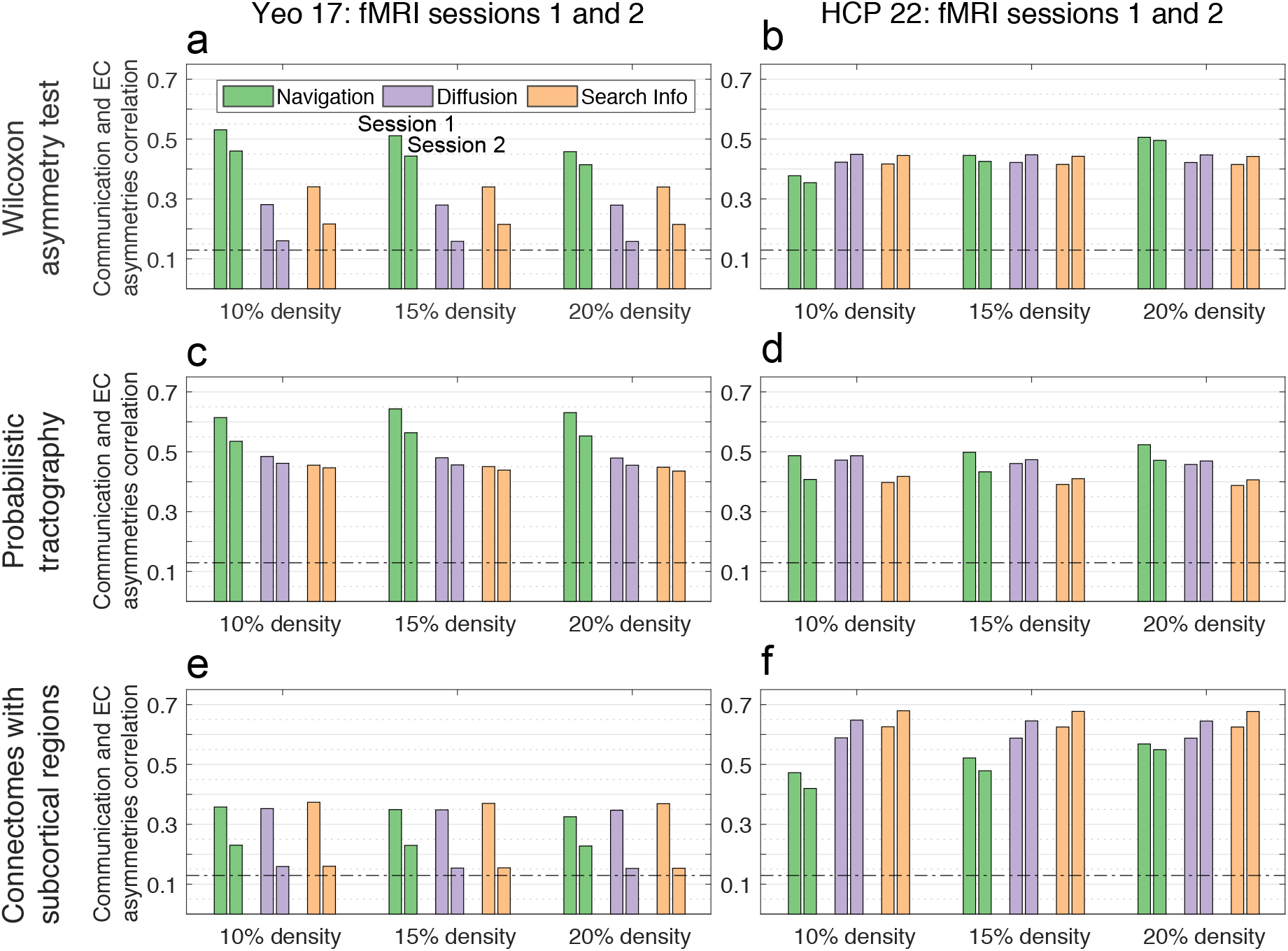
Relationship between send-receive asymmetry and directionality of effective connectivity across a range of methodological settings (*N* = 360). Bars denote the Pearson correlation coefficients between send-receive and effective connectivity asymmetries for *M* = 17 cortical subsystems. Bars are colored according to the three communication measures: i) navigation (green), ii) diffusion (violet), and iii) search information (beige). Correlations were computed for two independent resting-state fMRI sessions (Sessions 1 and 2) and multiple structural connection density thresholds (10, 15 and 20%). Significance threshold of *P* < 0.05 is indicated with a dotted line. **(a,b)** Non-parametric definition of send-receive asymmetry for *M* = 17, 22 cortical subsystems. **(c,d)** Send-receive asymmetry computed on connectomes derived using probabilistic tractography for *M* = 17, 22 cortical subsystems. **(e,f)** Send-receive asymmetry computed on connectomes including subcortical regions for *M* = 17, 22 cortical subsystems.

**Supplementary Fig. 11.**
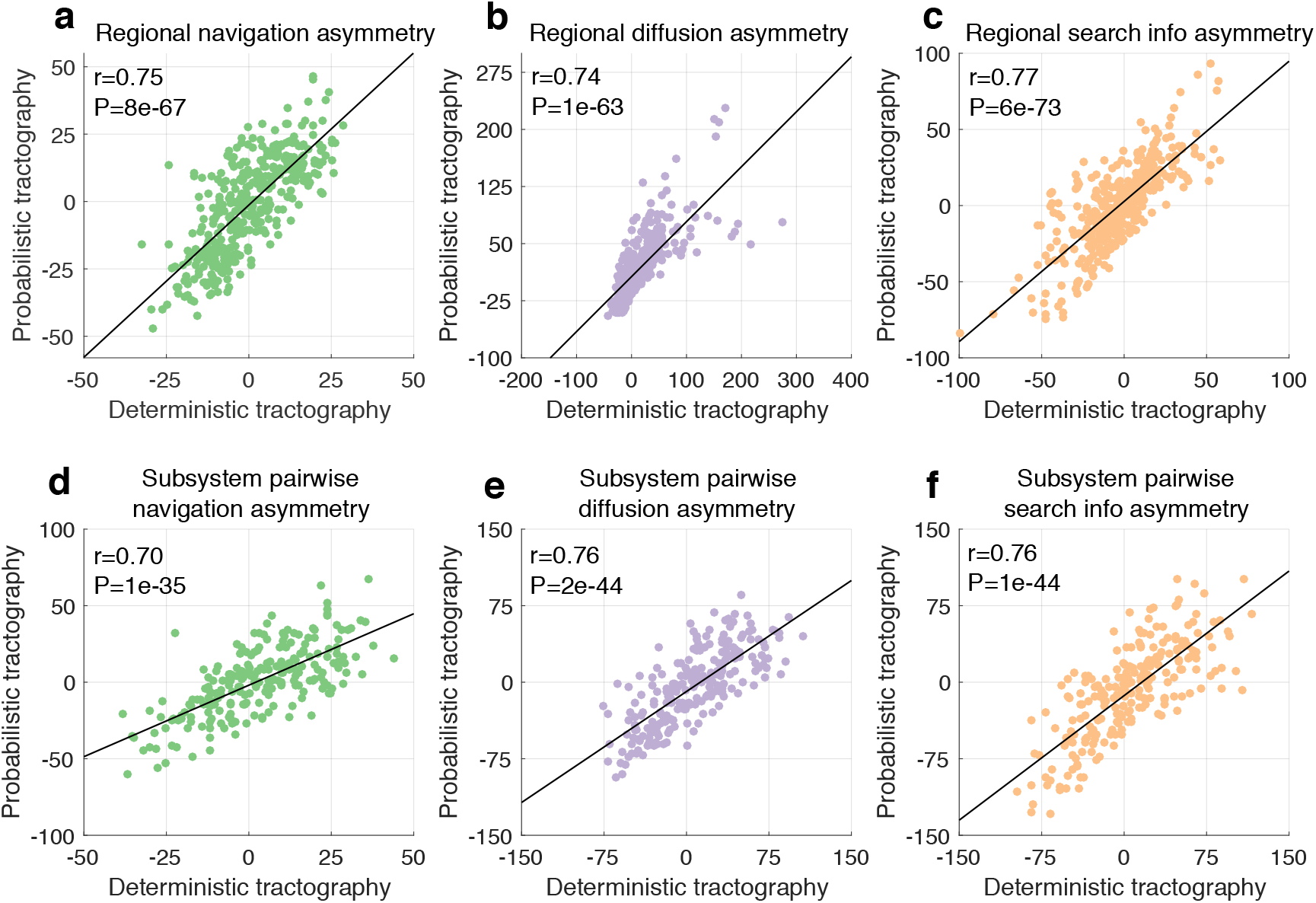
Scatter plots showing the relationship between send-receive asymmetries derived from deterministic and probabilistic connectomes (*N* = 360). **(a,b,c)** Deterministic and probabilistic regional send-receive asymmetries for navigation, diffusion and search information, respectively. **(d,e,f)** Subsystem pairwise send-receive asymmetry (*M* = 22) for navigation, diffusion and search information, respectively. *r*: Pearson correlation coefficient, *P*: associated P-value.

**Supplementary Fig. 12.**
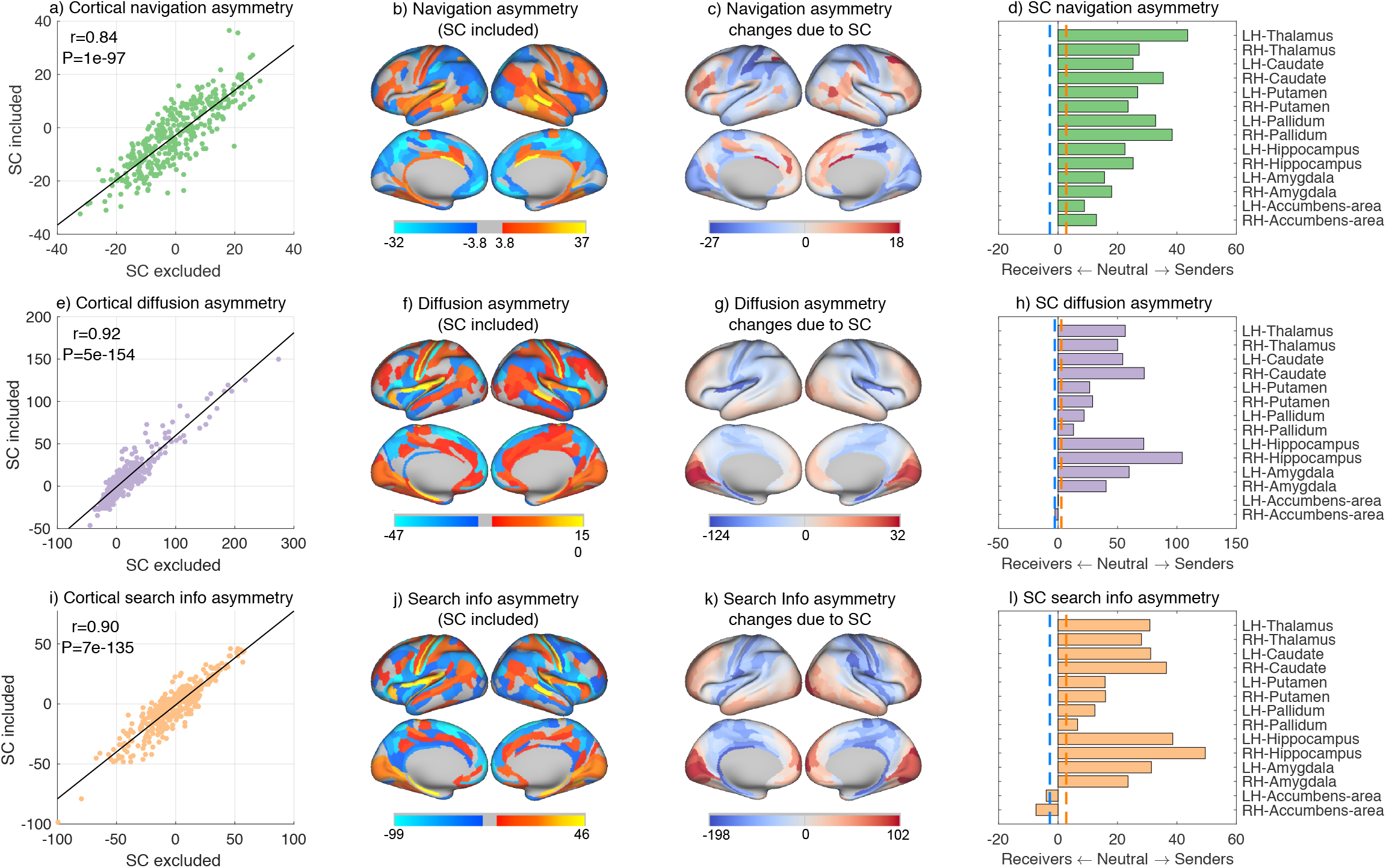
Regional send-receive asymmetry of human connectomes including subcortical regions. **(a)** Scatter plot of cortical navigation asymmetries computed from connectomes excluding subcortical structures (horizontal axis) and including subcortical structures (vertical axis). *r*: Pearson correlation coefficient, *P*: associated P-value. **(b)** Cortical navigation asymmetry computed from connectome including subcortical structures. Regions associated with significant send-receive asymmetry are classified as putative senders (orange) and receivers (blue). Regions colored gray are neutral and do not show significant send-receive asymmetry. **(c)** Differences in cortical navigation asymmetry following the inclusion of subcortical structures to the connectome. Positive differences (shown in red) indicate shifts towards outgoing communication efficiency, while negative differences (shown in blue) indicate shifts towards incoming communication efficiency. **(d)** Navigation asymmetry of subcortical regions. Dashed vertical lines indicate a significant bias towards outgoing (orange) and incoming (blue) communication efficiency. **(e–h)** Same as (a–d) for diffusion efficiency. **(i–l)** Same as (a–d) for search information.

**Supplementary Fig. 13.**
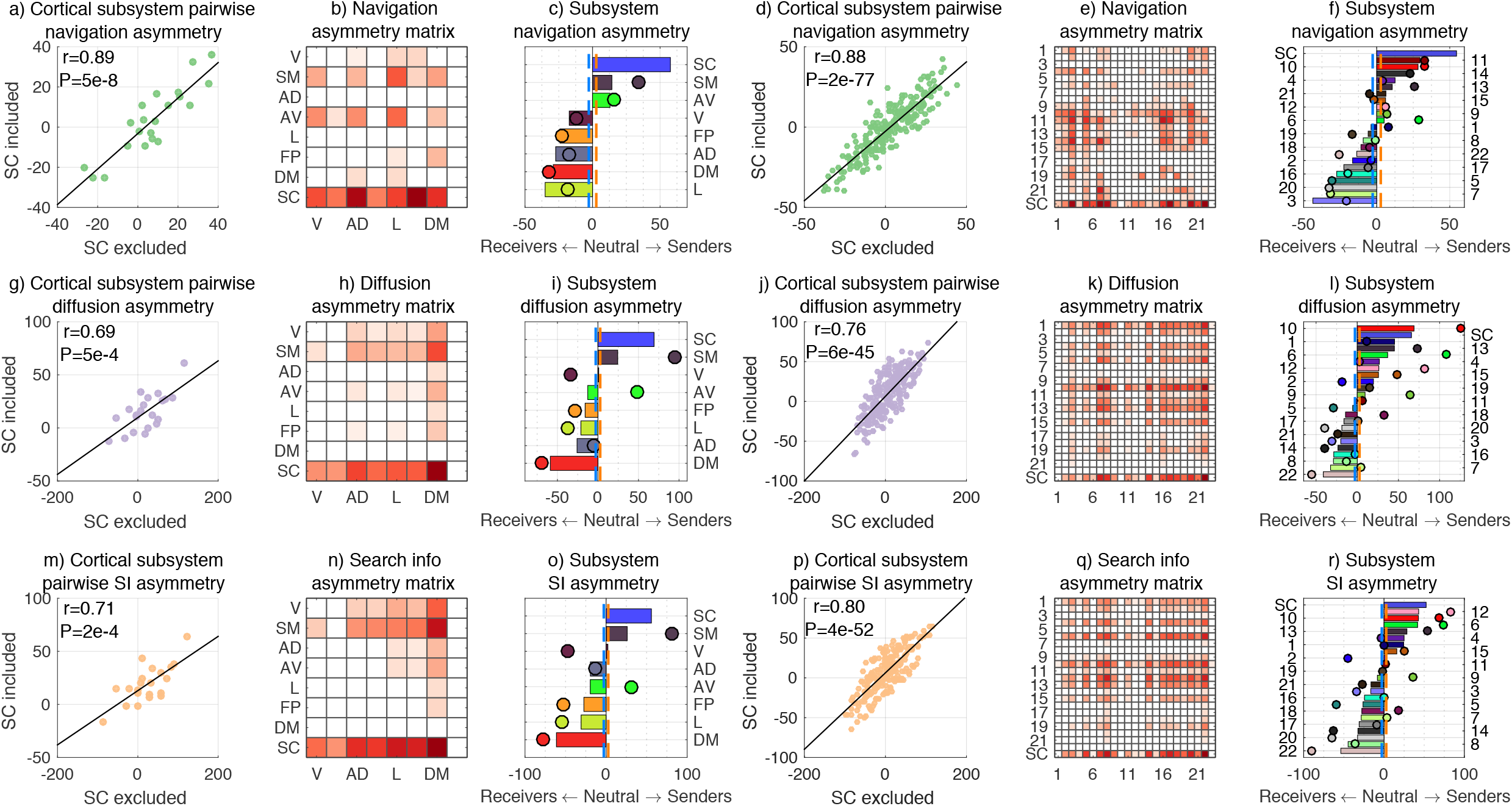
Pairwise subsystem send-receive asymmetry of human connectomes including subcortical regions. **(a)** Scatter plot of pairwise cortical subsystem (*M* = 7) navigation asymmetries computed from connectomes excluding subcortical structures (horizontal axis) and including subcortical structures (vertical axis). *r*: Pearson correlation coefficient, *P*: associated P-value. **(b)** Navigation asymmetry matrix from connectomes including the subcortex. Subcortical regions were grouped to form a single subsystem (SC), resulting in *M* = 7 + 1 subsystems. A matrix element *A*(*i, j*) > 0 denotes that communication occurs more efficiently from *i* to *j* than from *j* to *i*. Send-receive asymmetry values that did not survive multiple comparison correction were suppressed and appear as white cells. For ease of visualization and without loss of information (since *A*(*i, j*) = −*A*(*j, i*)), negative values were omitted. **(c)** Subsystems (*M* = 7 + 1) ranked by propensity to send (top) or receive (bottom) information under navigation. Send-receive asymmetries computed for connectomes including and excluding the subcortex are shown as bars and circles, respectively. Dashed vertical lines indicate a significant bias towards outgoing (orange) and incoming (blue) communication efficiency. **(d–f)** Same as (a–c) but for *M* = 22 + 1 subsystems. **(g–l)** Same as (a–f) for diffusion. **(g–l)** Same as (m–r) for search information.

**Supplementary Fig. 14.**
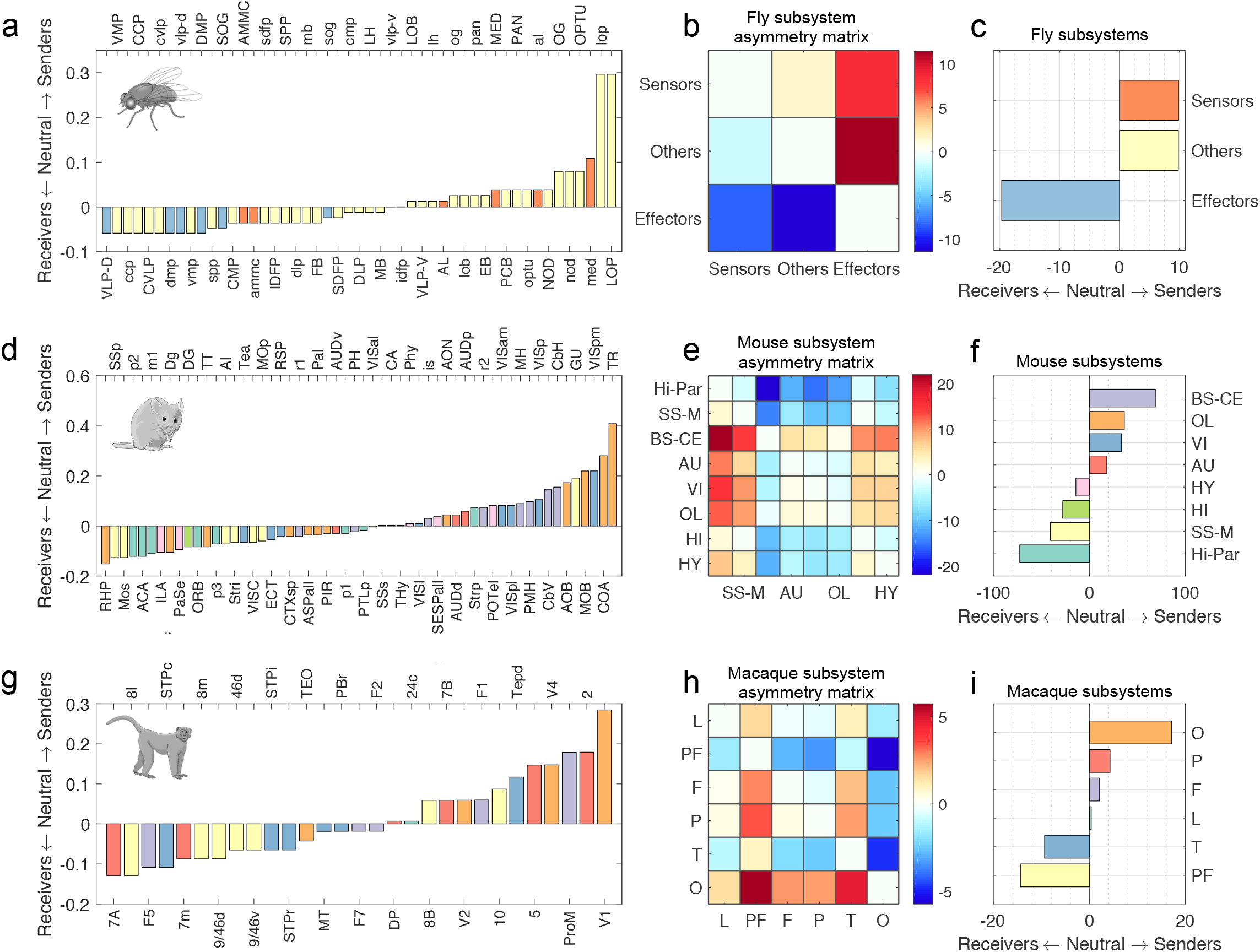
Send-receive asymmetry of undirected (symmetrized) non-human connectomes under binary diffusion efficiency. **(a)** Regions of the fly connectome sorted from receivers (negative send-receive asymmetry) to senders (positive send-receive asymmetry). Regions were divided into three subsystems [13], with sensor, effector and other nodes marked by orange, blue and beige bars, respectively. **(b)** Send-receive asymmetry matrix between subsystems of the fly connectome. For each subsystem pair *i, j*, we computed whether the mean of their distribution of node-level pairwise asymmetries was significantly larger than 0 by means of a one-sample t-test. Send-receive asymmetry between subsystems was defined as the resulting t-statistic. Red (positive t-statistic) and blue (negative t-statistic) indicate a bias towards communication efficiency in the *i* → *j* and *j* → *i* directions, respectively. **(c)** Fly subsystems sorted according to their send-receive asymmetry. Positive and negative values denote biases towards outgoing and incoming communication, respectively. **(d–f)** Same as (a–c) for the mouse connectome. Bar colors denote the affiliation of regions to previously identified subsystems [14]. **(g–i)** Same as (a–c) for the macaque connectome. Bar colors denote the affiliation of regions to previously identified subsystems [15].

**Supplementary Fig. 15.**
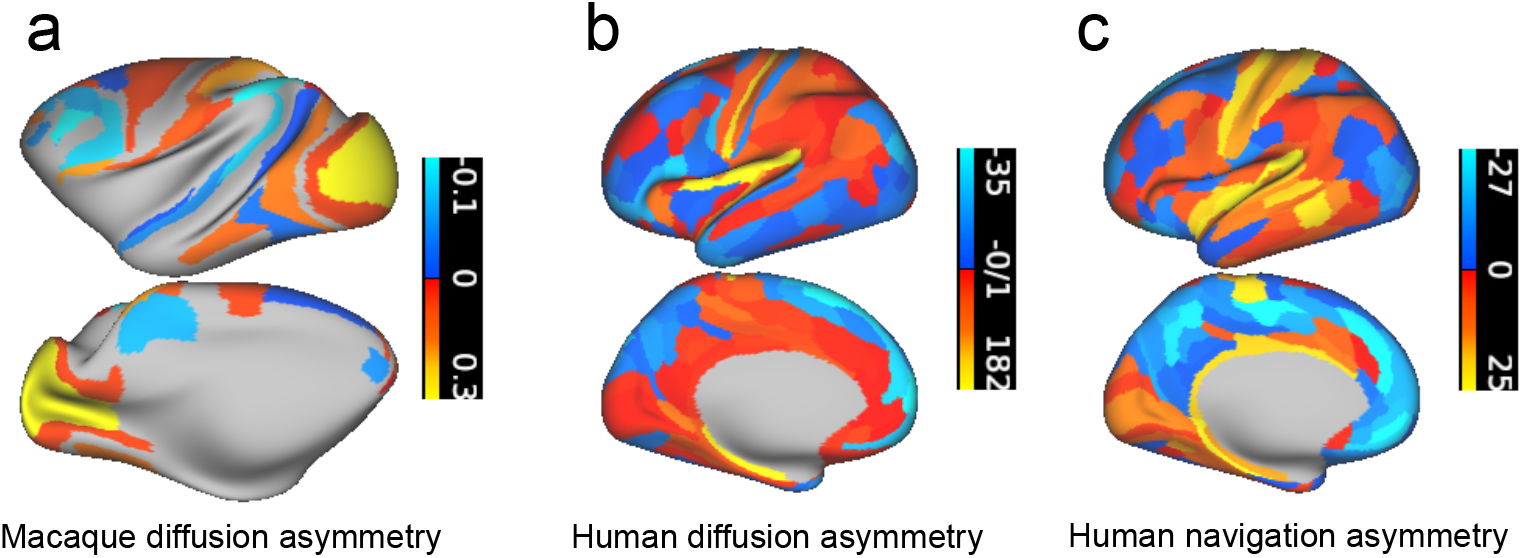
Comparison between the regional send-receive asymmetries for human and macaque. Orange and blue colors denote senders and receivers, respectively. **(a)** Binary diffusion asymmetry for the macaque. Gray regions are missing from the macaque connectome. **(b,c)** Weighted diffusion and navigation asymmetries for the human connectome.

